# A rise-to-threshold signal for a relative value deliberation

**DOI:** 10.1101/2021.09.23.461548

**Authors:** Vikram Vijayan, Fei Wang, Kaiyu Wang, Arun Chakravorty, Atsuko Adachi, Hessameddin Akhlaghpour, Barry J. Dickson, Gaby Maimon

## Abstract

Whereas progress has been made in identifying neural signals related to rapid, cued decisions^1–4^, less is known about how brains guide and terminate more ethologically relevant deliberations, where an animal’s own behavior governs the options experienced over minutes^5–8^. *Drosophila* search for many seconds to minutes for egg-laying sites with high relative value^9, 10^ and neurons, called *oviDNs*, exist whose activity fulfills necessity and sufficiency criteria for initiating the egg-deposition motor program^11^. Here we show that oviDNs express a calcium signal that rises over seconds to minutes as a fly deliberates whether to lay an egg. The calcium signal dips when an egg is internally prepared (ovulated), rises at a rate related to the relative value of the current substrate being experienced, and reaches a consistent peak just prior to the abdomen bend for egg deposition. We provide perturbational evidence that the egg-deposition motor program is initiated once this signal hits a threshold and that sub-threshold variation in the signal regulates the time spent deliberating and, ultimately, the option chosen. These results argue that a rise-to-threshold signal guides *Drosophila* to lay eggs on substrate options with high relative value, with each egg-laying event representing a self-paced decision similar to real-world decisions made by humans and other mammals.

---

Egg-laying site selection is critical for the survival of a fly’s progeny^12^. As such, *Drosophila* search for a high quality substrate for many seconds to minutes prior to depositing each individual egg^9, 10^. Over the past several years, egg-laying preferences or aversions for substrates varying along many dimensions have been documented, like stiffness^13^, illumination levels^14^, microbe content^15, 16^, and the concentrations of sucrose^9, 10^, ethanol^17, 18^, acetic acid^19^, polyamines^20^, terpenes (citrus compounds)^21^, and lobeline (a bitter compound)^22^. Despite the thorough work on the sensory side, how decision-related neural signals evolve in real time to guide the site selection process, and generate the observed preferences, is unknown. We sought to identify a neural signal that would help to explain how flies make adaptive decisions regarding where to lay their eggs in their local environment.

## A behavioral sequence for egg laying

We took videos of gravid *Drosophila* in a small chamber with a soft substrate floor. Using these videos, and building on past efforts^9, 11, 23–25^, we characterized a behavioral sequence for egg laying (see Supplementary Tables 1 and 2 for genotypes and conditions for all experiments). This characterization helped to define ethologically meaningful steps relevant to the neural measurements we studied later. The sequence begins with the fly standing still and performing an *abdomen elongation* and *scrunch* (Fig. 1a, steps 1 and 2), which likely relates to the internal act of *ovulation*^26^, i.e. passing an egg from an ovary to the uterus (see below). The fly then increases its locomotor speed (*search* period, step 3), performs an abdomen bend for egg deposition (step 4), deposits an egg (step 5), and performs an additional abdomen bend, likely for cleaning the ovipositor (step 6).

**Fig. 1.**
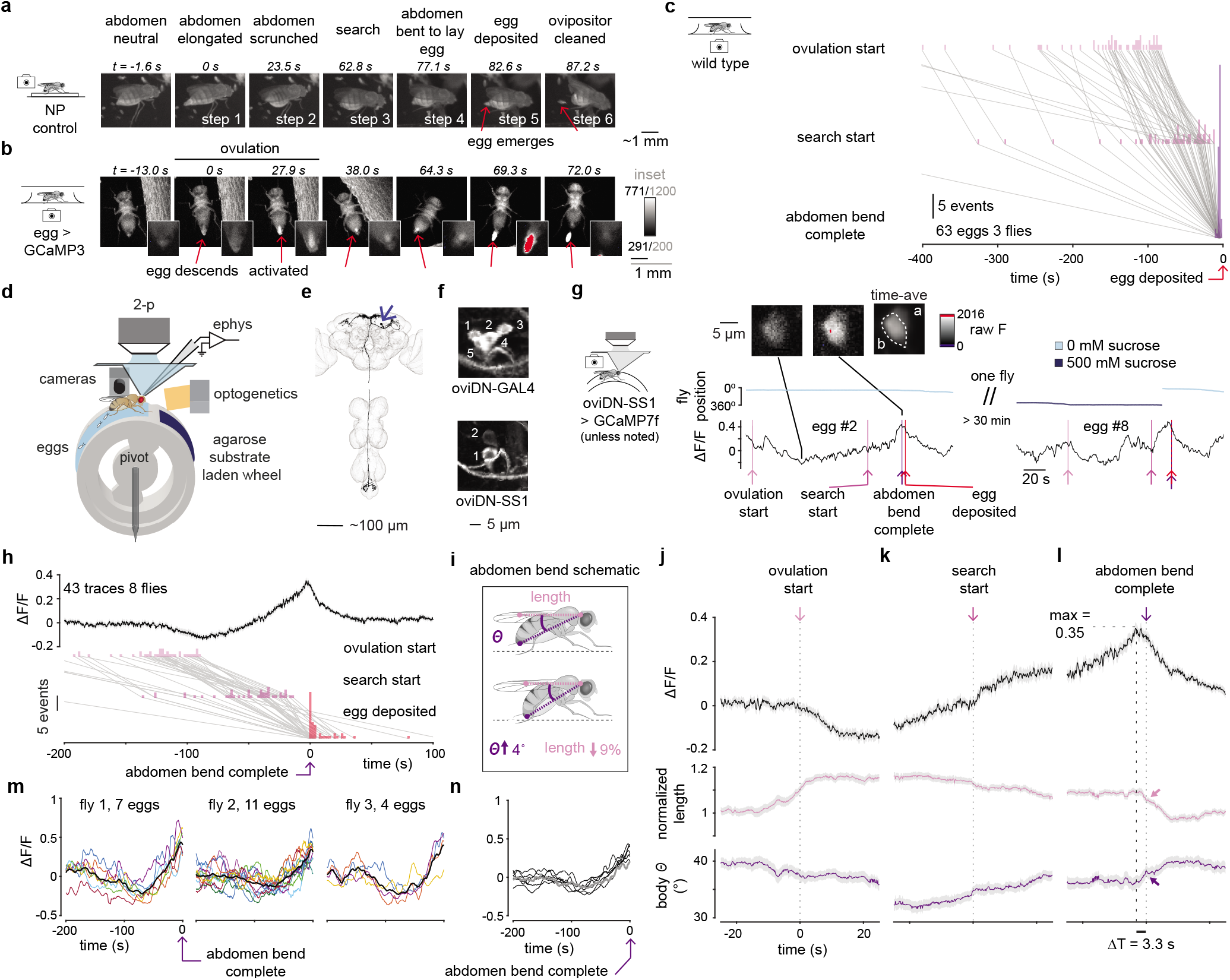
oviDN [Ca^2+^] dips during ovulation, rises for seconds to minutes, and peaks immediately before the abdomen bend for egg deposition. **a,** Behavioral sequence of egg laying. **b,** Imaging an egg expressing GCaMP3 in the body, over time. Steps correspond to panel a. Image contrast has been adjusted for display; insets show close ups, with over/under saturated pixels in red/blue. **c,** Progression of the behavioral sequence of egg laying. Grey lines connect events from a single egg-laying sequence in a single fly. **d,** Schematic of wheel for measuring neural activity and body posture during egg laying. **e,** A single oviDNb descending neuron traced from light-microscopy images. Arrow points to cell body. **f,** Images of oviDN and oviDN-like cell bodies in a single side of the brain labeled by the two driver lines used in this study. The fly’s midline is on the left. **g,** oviDN ΔF/F and fly behavior during egg laying for two eggs laid by the same fly. ΔF/F is smoothed with a 2 s boxcar filter. Images show the z-projection of selected two-photon imaging slices. Labels a and b in images refer to oviDNa and oviDNb, respectively (oviDNa is partially obscured by oviDNb in the z-projection). **h,** Population-averaged oviDNb ΔF/F aligned to the end of the abdomen bend for egg laying (light gray shading is s.e.). Data associated with 43 imaging traces from 41 egg-laying events associated with 9 cells in 8 flies are shown. The number of traces exceeds the number of egg laying events by two because for two eggs we imaged the oviDNb on both sides of the brain. Behavioral events shown below. **i,** A schematic of abdomen bend. θ is body angle and length is neck to ovipositor distance. **j-l,** Mean oviDN ΔF/F and fly behavior aligned to events in behavioral sequence shown in panel h. Normalized length is the length as shown in panel i divided by the median length (Methods). Filled arrows point to the moment the abdomen bend for egg deposition is complete. A subsequent (stronger) bend is, presumably, for cleaning the ovipositor after egg deposition. **m,** oviDN ΔF/F during individual egg-laying events for 3 separate flies, smoothed with a 5 s boxcar filter. Mean in black. **n,** ean oviDN ΔF/F during egg laying for all 7 flies that laid ≥ 3 eggs, smoothed with a 5 s boxcar filter. A single fly imaged with GCaMP7b is shown with a thicker grey line.

This behavioral sequence is consistent with those described previously^9, 11, 23–25^ and although abdomen movements prior to egg laying have been noted^23–25^, they have not been definitively linked to ovulation. To directly test whether steps 1 and 2 (abdomen elongation and scrunch) reflect ovulation, we expressed GCaMP^27^ in eggs^28^, which allowed us to visualize eggs moving from the ovaries to the uterus inside the fly. We could also visualize when each egg was activated so as to start embryonic development because egg activation is associated with a large [Ca^2+^] increase inside the egg^28^, yielding a boost in the GCaMP signal. We imaged flies from below in custom, high-resolution egg-laying chambers with soft, agarose floors that promote egg laying (Extended Data Fig. 1a) (Methods). We observed that an egg indeed descends from an ovary to the uterus during abdomen elongation (Fig. 1b, step 1) and the egg furthermore exhibits a strong increase in GCaMP fluorescence during abdomen scrunching (Fig. 1b, step 2) (Supplementary Video 1), confirming that the elongation and scrunch reflect ovulation.

We quantified the egg-laying behavioral sequence in wild type flies (Methods). Specifically, for each egg we annotated four (of the six) egg-laying steps just mentioned: (1) *ovulation start* (when the abdomen first begins to elongate), (2) *search start* (when the abdomen returns to a neutral posture after ovulation), (3) *abdomen bend complete* (the time point of the maximum abdomen deflection prior to egg deposition), and (4) *egg deposition* (when half the egg is visible outside the ovipositor) (Fig. 1c, Extended Data Fig. 2). Might there exist a neural signal that explains variability in this behavioral sequence, in particular the variability between the *search start* time and the *abdomen bend for egg deposition*, that is, how long the fly spends searching for a good substrate?

## A preparation for neurophysiology during egg laying

We developed an agarose-laden cylindrical treadmill––i.e. an egg-laying *wheel*––that a head-fixed fly can rotate as it walks, and lay eggs on, while we perform two-photon imaging or electrophysiological recording from neurons in the brain (Fig. 1d, Extended Data Fig. 3) (Methods). Each wheel had regions for agarose, with thin plastic barriers in-between, and the agarose substrates could vary in their sucrose concentration (Fig. 1d, e.g., light and dark blue). All agarose substrates––in these head-fixed experiments as well as in later free-behavior experiments––contained 1.6% ethanol and 0.8% acetic acid, which simulates the environment of a rotting fruit and promotes egg laying. We found that the egg-laying behavioral sequence measured on the wheel resembled that in free behavior (Extended Data Fig. 4a-b).

However, one difference was that flies walked less vigorously during the *search* period while on the wheel (compare the speed of the flies in Extended Data Fig. 4f and Extended Data Fig. 2c). Reduced locomotion during head-fixed search likely results from the fact that flies find it difficult to start rotating the wheel from rest, due to its inertia, and the wheel is necessarily stationary after the extended pause in locomotion associated with ovulation (Methods). As such, we consider the *search* period as a *search/delay* period on the wheel.

A recent study identified oviposition descending neurons (*oviDNs*) in *Drosophila*^11^. These neurons are key for egg laying because when they are inhibited egg laying is completely suppressed and when they are stimulated an egg is often laid^11^. Three oviDNs^11^ and two uncharacterized oviDN-like neurons, are present in a single side of the female hemibrain electron-microscopy volume^29^ (totaling ten neurons per brain) (Extended Data Fig. 5a). The two uncharacterized oviDN-like neurons per side have similar morphology to the characterized oviDNs with each neuron primarily receiving input in the central brain and primarily sending output to the abdominal ganglion (Fig. 1e). We used two different driver lines to gain genetic access to oviDNs––VT040574 (hereafter, oviDN-GAL4, Extended Data Fig. 5b) and oviDN-SS1^11^. OviDN-GAL4 labels 3 of 3 oviDNs and 2 of 2 oviDN-like neurons per side and oviDN-SS1 labels 2 of 3 oviDNs (cholinergic neurons named oviDNa and oviDNb)^11^ and 0 of 2 oviDN-like neurons per side (Fig. 1f, Extended Data Fig. 5a). For brevity, we hereafter refer to the group of ten neurons as oviDNs. For two-photon imaging, we used the oviDN-SS1 driver and targeted the cell body of oviDNb (unless otherwise stated), which has the brighter GCaMP^30^ signal of the two neurons (Extended Data Fig. 6a). Rather than targeting intermixed neurites, we typically targeted the lone oviDNb soma on one side, so that we could consistently image the same identified cell from all flies.

## oviDN [Ca^2+^] dips during ovulation, rises for seconds to minutes, and peaks immediately before the abdomen bend for egg deposition

We imaged GCaMP7 fluorescence in oviDNs during egg laying (Fig. 1g-l). We found that the oviDN ΔF/F signal drops to its minimum value during ovulation and then generally rises until the moment when the abdomen bends to deposit an egg (Fig. 1g). In some cases, we observed a monotonic rise (Fig. 1g left, Supplementary Video 3). In other cases, the signal fluctuated prior to reaching its peak (Fig. 1g right, Supplementary Video 4). The peak in the population-averaged ΔF/F signal was higher when we aligned the oviDN [Ca^2+^] signal to the moment when the abdomen finishes bending to lay the egg (Fig. 1h-i) rather than aligning to the moment when the egg becomes half visible outside the fly (Extended Data Fig. 4g vs. Extended Data Fig. 4h). On average, the [Ca^2+^] signal dips when ovulation starts (Fig. 1j), reaching its minimum when the abdomen is longest (Extended Data Fig. 4d). We interpret this signal dip as a ‘zeroing event’ that occurs with ovulation. The average [Ca^2+^] signal then begins to rise and returns to near baseline (ΔF/F of 0 in our normalization) (Methods) when ovulation is complete, which we define as the beginning of the search/delay period (Fig. 1k). We often observed in individual traces an upward inflection in the [Ca^2+^] rise right after the search/delay period is initiated (Fig. 1g right, after the search-start arrow) which is evident as a slight inflection in the mean trace (Fig. 1k, after the search-start arrow). The average [Ca^2+^] signal peaks ∼3 s prior to when the abdominal bend to deposit an egg is complete (Fig. 1l), that is, approximately when the bend is initiated. The average [Ca^2+^] signal returns to baseline after egg laying, as flies perform an additional abdomen bend presumably associated with grooming their ovipositor (Extended Data Fig. 4i).

The [Ca^2+^] rise was present across multiple egg-laying events from individual flies (Fig. 1m), reaching a qualitatively similar ΔF/F value of ∼0.35 immediately prior to the abdomen bend for egg laying (Fig. 1n). A ΔF/F of 0.35 means that the GCaMP signal is 35% higher than at baseline, with the baseline calculated as the mean fluorescence in the surrounding 20-minute window (Methods). A relatively modest, 35%, signal rise makes sense since cells are not thought to be capable of sustaining strongly elevated calcium levels for many minutes^31^. In some flies, we simultaneously imaged additional cells, like the ipsilateral oviDNa or contralateral oviDNb; the oviDNa showed a similar rising signal, which also peaked just prior to the abdomen bend for egg deposition (Extended Data Fig. 6b). The GCaMP signal from both the ipsilateral oviDNa and the contralateral oviDNb showed a cross-correlation with a zero-lag peak with the oviDNb (Extended Data Fig. 6c-d), supporting a model where all four oviDNs in the oviDN-SS1 line exhibit the same first-order calcium dynamics during egg laying. During non-egg-laying periods the oviDN ΔF/F signal fluctuated, maintaining a correlation with abdomen movements and locomotor variables (Extended Data Fig. 7a-d). Approximately once every half hour, the oviDN ΔF/F signal reached the 0.35 level without ovulation having occurred prior and these moments were associated with abdomen bends that yielded no egg (Extended Data Fig. 7e).

oviDNs thus express a signal whose dynamics correlate tightly with the behavioral sequence of *Drosophila* egg laying. This discovery raises two key questions that we pursue further here. First, is the egg-deposition motor program triggered by oviDN activity hitting a threshold? Second, if an activity threshold indeed triggers egg deposition, is subthreshold oviDN activity modulated during the search period to increase the chance that threshold is reached on substrates of higher relative value?

## Evidence for a threshold in oviDN [Ca^2+^] activity triggering the egg-deposition motor program

To test whether the egg-deposition motor program is initiated when oviDNs hit a threshold, we co-expressed in them GCaMP7f and the light-gated ion channel, CsChrimson^32^. We then measured oviDN ΔF/F and fly behavior while providing 5 s long, high-intensity light pulses (Methods). Stimulations after ovulation typically yielded an abdomen bend and egg deposition (Fig. 2a, Supplementary Video 5). Of the 34 high-intensity stimulations after ovulation, 28 (82%) resulted in egg deposition. Of the 174 high-intensity stimulations without a prior ovulation event, 4 (2%) resulted in eggs (Fig. 2b). In all 4 of those cases, the eggs were from the first stimulation pulse given to a fly and ovulation may have occurred prior to the imaging session. If we average [Ca^2+^] and behavioral signals around the time of stimulations that yielded an egg, we observe an increase in ΔF/F in the oviDN alongside a synchronous abdomen bend and egg deposition with variable latency (Fig. 2c). Note that the peak in oviDN [Ca^2+^] slightly lagged the initiation moment of the abdomen bend in these stimulation experiments. It is likely that with optogenetic light pulses, however, that [Ca^2+^] at presynaptic zones––the relevant compartment for driving behavior–– reaches threshold earlier than does [Ca^2+^] at the soma, given that synapses are small and are expected to have faster [Ca^2+^] dynamics than cell bodies^33^.

**Fig. 2.**
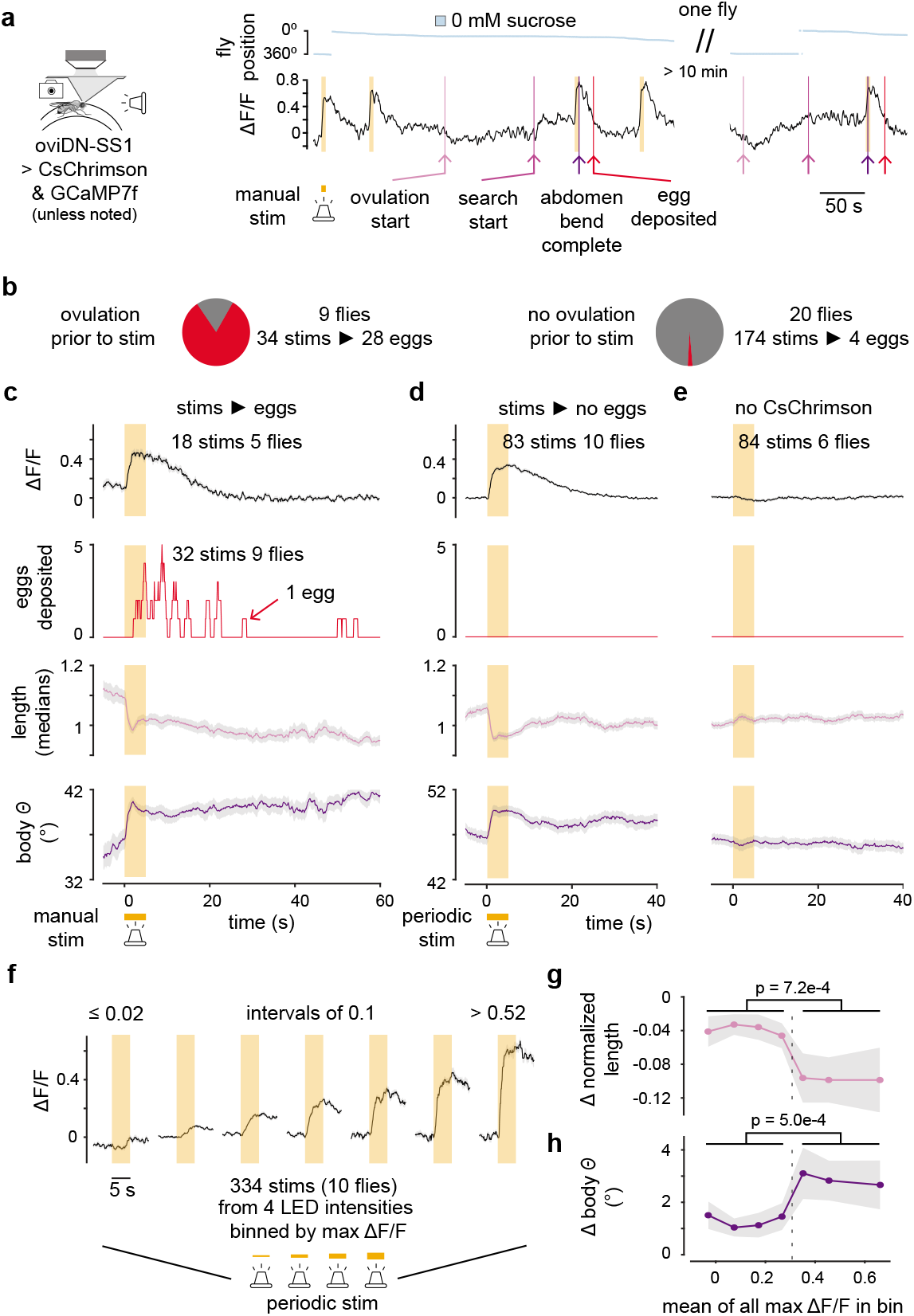
Evidence for a threshold in oviDN [Ca^2+^] activity triggering the egg-deposition motor program. **a,** Sample trace of an oviDN ΔF/F signal and behavior during high-intensity 5 s CsChrimson stimulation (stim.). ΔF/F is smoothed with a 2 s boxcar filter. **b,** Pie charts indicating the fraction of high-intensity CsChrimson stimulations that resulted in egg deposition within the subsequent 60 s, separated based on whether ovulation was or was not observed prior. **c,** ean oviDN ΔF/F and behavior for manually triggered high-intensity 5 s CsChrimson stimulations that resulted in egg deposition within the subsequent 60 s. For behavior: 32 stimulations in 9 flies. For ΔF/F: 18 stimulations in 5 flies. (There are more traces contributing to the behavioral average than the ΔF/F average because bleed-through of optogenetic stimulation light into our imaging PMTs contaminated initial ΔF/F measurements) (Methods). **d,** Mean oviDN ΔF/F and behavior for periodically triggered high-intensity 5 s CsChrimson stimulations that did not result in egg deposition within the subsequent 60 s. **e,** Same as panel d but with flies not expressing CsChrimson. **f,** oviDN ΔF/F responses to optogenetic stimulation binned by the max ΔF/F 1 to 3 s after stimulation starts. Stimulations at four different intensities were triggered periodically, which generated a range of oviDN ΔF/F responses. **g, h,** Change in mean body length and body angle for each of the bins shown in panel f. The mean 2 to 4 s after stimulation starts was subtracted from the mean 0 to 2 s prior to stimulation.

In our initial experiments, we stimulated the oviDNs at user-defined moments. In subsequent experiments, we performed regularly spaced stimulations in flies expressing or not expressing CsChrimson. Flies expressing CsChrimson bent their abdomen, on average, with regularly spaced stimulation, even as none of these stimulations resulted in an egg (Fig. 2d), whereas control flies did not bend their abdomen (Fig. 2e). We interpret this result––alongside the observation that flies tend to bend their abdomen when oviDN activity is spontaneously high without prior ovulation (Extended Data Fig. 7e)––to mean that flies initiate the egg-deposition motor program when oviDNs reach a high level of activity. If an egg is available in the uterus, egg deposition occurs, albeit with temporal variability that is perhaps related to sensory feedback signals in the uterus^24^ or aspects of the motor sequence related to how eggs are released^25^. The temporal variability in egg deposition was qualitatively similar in optogenetically-stimulated (Fig. 2c) and spontaneous (Fig. 1h) egg laying in head-fixed flies.

Is the egg-deposition motor program initiated in an all-or-nothing fashion when oviDN activity crosses a threshold? To test this hypothesis, we stimulated oviDNs at a regular interval while cycling through four different intensities of light. We assigned each oviDN response to one of seven bins depending on the ΔF/F maximum on that stimulation pulse (Fig. 2f). For each bin, we also plotted how much the abdomen moved. We found that when the stimulation pulse caused the oviDN ΔF/F signal to exceed ∼0.35, the flies produced a large abdomen bend and when the stimulation pulse induced a smaller oviDN signal, there was not a large bend (Fig. 2g-h). This result was robust to how we binned ΔF/F responses (Extended Data Fig. 8). These data are consistent with the hypothesis that a threshold level of oviDN activity initiates the egg-deposition motor program in a yes/no fashion. It is intriguing that the threshold level for abdomen bending in these CsChrimson stimulation experiments is similar to the 0.35 ΔF/F value observed during spontaneous egg laying (Fig. 1l). Because the excitability of CsChrimson expressing neurons is likely to be different from neurons lacking an optogenetic opsin and we are only stimulating 2 of the 5 oviDNs per side, this correspondence is not guaranteed to be functionally meaningful.

## Flies search for an egg-laying substrate with high relative value in the time period when the oviDN [Ca^2+^] is rising

If an oviDN activity-threshold triggers initiation of the egg-deposition motor program, might substrate quality modulate oviDN activity to influence when threshold is reached and thus where an egg is laid? We first analyzed the egg-laying behavior of freely walking *Drosophila* to better understand how flies use substrate experiences during the search period, i.e., the time period after ovulation and before egg-deposition is initiated, to guide egg-laying substrate choice. We placed flies in custom, high-throughput chambers for egg-laying with wells for two different substrate options (Fig. 3a, Extended Data Fig. 1b, Supplementary Video 6) (Methods). We focused on sucrose-based choice because sucrose is non-volatile and this allowed us to infer the timescale over which flies use past substrate experiences to guide egg laying on the current option, independently of the flies being able to smell, or otherwise sense, the previous option at a distance (see below).

**Fig. 3.**
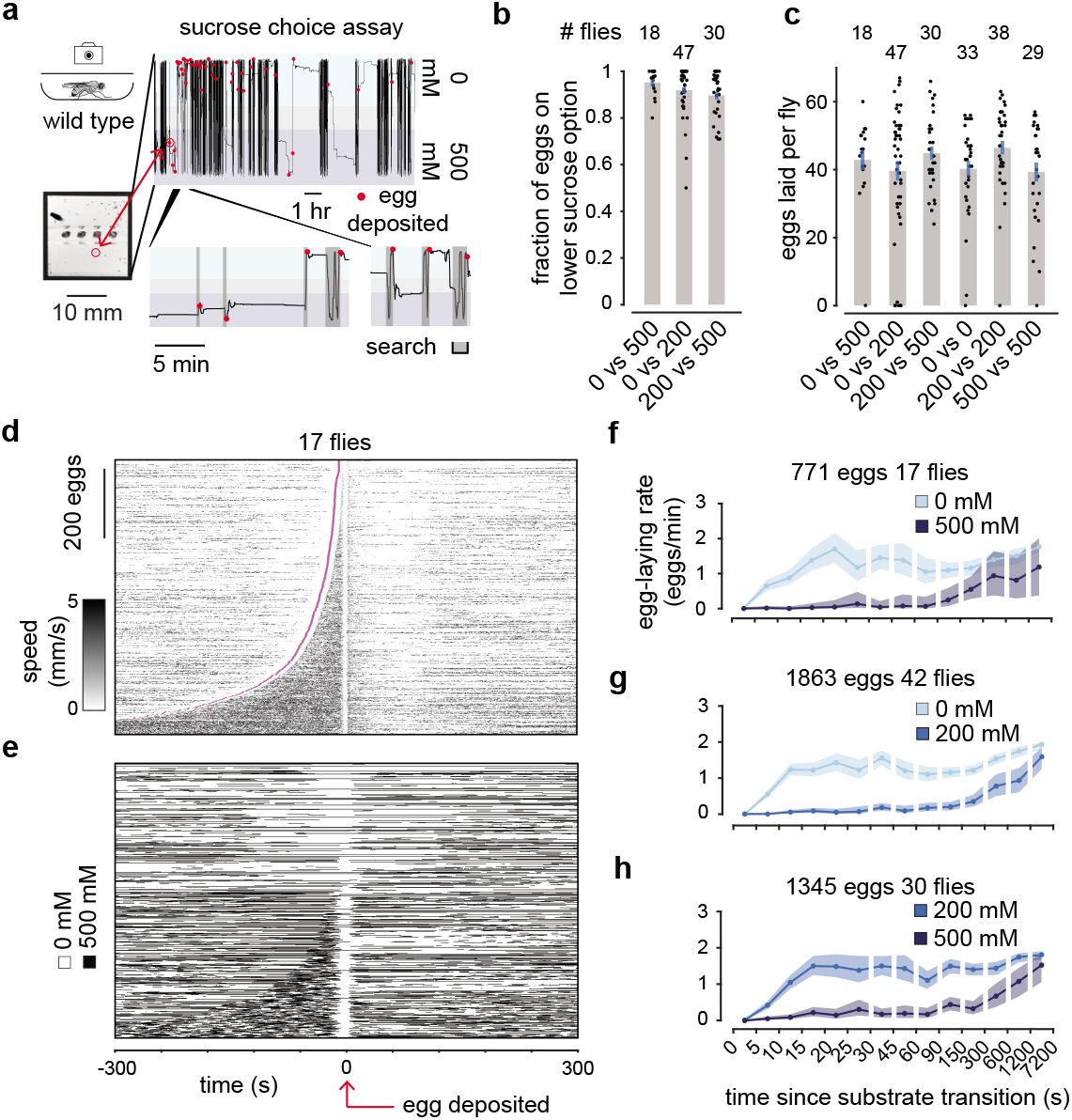
Flies search for a high relative value egg-deposition site during the period the oviDN [Ca^2+^] rises. **a,** Y-position trajectory, and egg-deposition events from a single fly in a high-throughput egg-laying choice chamber. **b,** Fraction of eggs on the lower sucrose option with 95% confidence interval. x-axis indicates sucrose concentration in mM. One dot is one fly. **c,** Eggs laid per fly. One dot is one fly. **d,** Each row represents a single egg-laying event in a 0 vs. 500 mM sucrose chamber, aligned to egg deposition, with the fly’s speed indicated by the black intensity. Rows have been ordered based on the search duration. Start of the search period is indicated in magenta. 18 flies were tested, and one did not lay eggs. **e,** Same data as in panel d, but the substrate, 0 vs. 500 mM, that the fly was residing on is indicated with white and black pixels respectively. **f-h,** Mean egg-laying rate during the search period after a transition from higher to lower sucrose (lighter blues) or lower to higher sucrose (darker blues) in three separate sucrose choice conditions with 90% confidence interval shaded (Methods). 771 eggs from 17 flies (18 flies tested and 1 did not lay eggs), 1863 eggs from 42 flies (47 flies tested and 5 did not lay eggs), and 1345 eggs from 30 flies (30 flies tested), respectively.

*Drosophila melanogaster*, perhaps counterintuitively, lay most of their eggs on the substrate with the lower, not higher, concentration of sucrose^9, 10^. In the wild, *D. melanogaster* prefer to lay eggs on rotting fruit^13^, and the bias to low sucrose in our assays may be because a soft substrate with high levels of ethanol and low levels of sucrose^16^ mimics the rotting portion of the fruit where active fermentation is taking place (i.e., conversion of sugar to alcohol). We replicated past results that showed sucrose-based choice to be, remarkably, a relative-value decision^9, 10^. That is, flies strongly bias egg laying to the lower of two sucrose options rather than preferring an absolute sucrose concentration. For example, they laid > 90% of eggs on the 0 mM option in 0 vs. 200 mM chambers and they laid > 90% of their eggs on the 200 mM option (the previously avoided substrate) in 200 vs. 500 mM chambers (Fig. 3b). Flies laid a similar total number of eggs in all chambers, regardless of whether the chambers had substrates with the same sucrose concentration in both wells (one option) or two different concentrations (two options)^9, 10^ (Fig. 3c).

Although we did not collect images with sufficient spatial resolution to detect elongation and scrunching of the flies’ abdomens in these chambers, we could still detect the onset of the search period by analyzing the locomotor trajectories of flies because *Drosophila* consistently stand still for ∼1 min. when they ovulate (Extended Data Fig. 2b-c) (Methods). We could also denote the end of the search period as the moment at which an egg was half out of the ovipositor, which follows the final abdomen bend for egg laying by only a few seconds in these environments (Fig. 1c) (Fig. 3a) (Methods). The duration of the search period was highly variable (Fig. 3d). The search almost always ended with flies laying an egg on the lower sucrose option, despite spending appreciable time on the higher option during the search epoch^10^ (Fig. 3e). 95% (734 of 771) of eggs were laid on 0 mM while only 77% (592 of 771) of search periods started on 0 mM (p = 2.1e-25). Note that more search periods start on 0 mM because ovulation tends to occur soon after the previous egg-laying event (Extended Data Fig. 2a) and egg laying tends to occur on 0 mM. We also noticed that when flies started the search period on 500 mM, they frequently left the high-sucrose substrate while searching (83%, 149 of 179), but when flies started their search on 0 mM, they left the low-sucrose substrate less often (36%, 212 of 592) (p = 8.4e-29). Leaving a higher sucrose substrate more often at the onset of search is not an intrinsic property of the substrate because flies leave substrate islands at a similar rate in 500 vs. 500 and 0 vs. 0 mM chambers (299 of 528 = 57% and 441 of 895 = 49%, respectively). Flies thus retain information about the substrate options available to them from experiences outside of the current search period and use this information to regulate the search. We tested for the possibility of flies using spatial memories to guide their egg-laying behavior in our chambers, but we could not find supportive evidence (Extended Data Fig. 9). Flies probably use spatial memories to guide egg-laying behavior in other contexts, but our experiments were conducted in darkness, in chambers where flies typically run around the chamber edges continuously (thigmotaxis), which likely promotes a non-spatial strategy. We also did not find evidence that flies were specifically pausing to feed on the higher sucrose substrate while searching, suggesting that in our experiments a competing feeding drive is not the reason for suppression of egg laying on higher sucrose substrates (Extended Data Fig. 9).

We noticed that flies would occasionally lay eggs on the higher sucrose option, specifically if a few minutes had elapsed since they last visited the preferred, lower sucrose option (Fig. 3a bottom, first two eggs). To quantify this observation, we calculated the egg-laying rate during the search period as a function of time since the last substrate transition (regardless of whether the last transition occurred in the current search period or prior) (Methods). Flies in 0 vs. 500 mM sucrose choice chambers strongly inhibited egg laying on 500 mM if they had visited the 0 mM option in the past ∼2 min. (Fig. 3f). After ∼2 min., the egg-laying rate on 500 mM began to increase gradually, approaching, albeit not completely, the egg-laying rate on 0 mM at the 2-hour time point. One interpretation of this egg-laying-rate plot is that the relative value of the 500 mM substrate gradually increases over time, eventually approaching the value of the 0 mM substrate, if 0 mM is not revisited. This phenomenon is also visible in 0 vs. 200 mM and 200 vs. 500 mM chambers (Fig. 3g-h). The fact that the egg-laying rate takes ∼10 s to reach its maximum value after a fly transitions to the higher relative value (lower sucrose) option (Fig. 3f-h) is due, at least in part, to the fact that flies do not lay eggs on the ∼2.5 mm plastic boundary between substrates (Extended Data Fig 10a-b) and also to the fact that there is a ∼3 s delay between when flies bend their abdomen to lay an egg and when the egg is deposited (Extended Data Fig 10c, Fig. 1c). Thus, the fly’s internal sense of relative value likely changes more rapidly after a transition than the slowly increasing egg-laying-rate curve would suggest.

## The relative value of the current egg-laying option influences the subthreshold physiology of oviDNs to impact when threshold is reached

How might oviDN physiology guide flies to lay eggs on the option with higher relative value? Perhaps the rate of rise in oviDN [Ca^2+^] during search is modulated by relative value (Fig. 4a)? If so, flies on a high relative-value substrate might show an oviDN [Ca^2+^] signal that rises faster and thus hits threshold more quickly. Flies on a low relative-value substrate would have their [Ca^2+^] rise more slowly, or even fall, affording the flies more time to find a better option before threshold is reached.

**Fig. 4.**
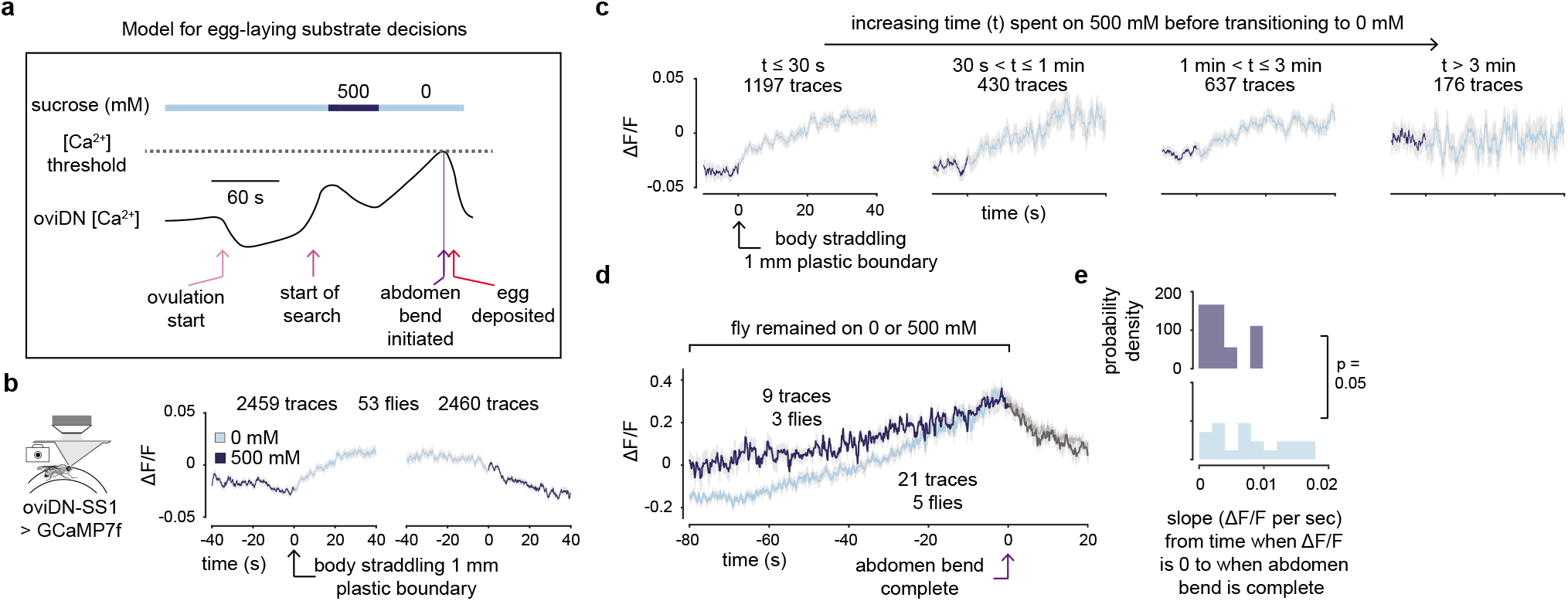
The relative value of the current egg-laying option influences the subthreshold physiology of oviDNs to impact when threshold is reached. **a,** Model for how the oviDN signal can underlie substrate decisions for egg-laying. **b,** Mean oviDN ΔF/F during substrate transitions from 500 to 0 mM and 0 to 500 mM. 2459 and 2460 traces from 70 cells in 53 flies (1911 and 1922 transitions). 1911 transitions yielded 2459 traces because we sometimes imaged the oviDNb cell on both sides of the brain. **c,** Mean oviDN ΔF/F during substrate transitions from 500 to 0 mM sucrose split based on the amount of time the fly spent on 500 prior to transitioning to 0 mM. 1197, 430, 637, and 176 traces from 70 cells in 53 flies (914, 347, 486, and 148 transitions). **d**, Mean oviDN ΔF/F for egg-laying events where the fly stayed on 0 mM (light blue) or 500 mM (dark blue) for the 80 s window prior to and including egg deposition. For 0 mM, 21 traces from 5 cells in 5 flies (21 eggs) are averaged. For 500 mM, 9 traces from 4 cells in 3 flies (7 eggs) are averaged. **e,** Probability densities of the individual oviDN ΔF/F slopes from the egg-laying events averaged in panel d. Individual oviDN ΔF/F’s were smoothed with a 5 s boxcar filter prior to calculating slopes.

We analyzed oviDN ΔF/F changes for all substrate transitions on the wheel. On wheels with 0 and 500 mM sucrose options, we observed an increase in the average ΔF/F after flies walked onto the higher relative-value substrate (decrease in sucrose concentration) and a decrease after flies transitioned to the lower relative-value substrate (increase in sucrose concentration) (Fig. 4b). Note that flies often straddled the boundary between substrates (Extended Data Fig. 11a), which will cause the change in [Ca^2+^] with substrate transitions to appear more gradual than it would be otherwise. The difference in ΔF/F on the two substrates was not explained by persistent differences in feeding or locomotor speed between the two options (Extended Data Fig. 11b-c). We observed similar changes in oviDN activity with substrate changes at the level of the oviDN membrane potential and spike rate (Extended Data Fig. 12).

If oviDN [Ca^2+^] tracks the relative value of substrates, rather than just sucrose concentration, one might expect that oviDN activity would gradually increase on the 500 mM option, as that option slowly becomes more acceptable over several minutes. Indeed, when we split 500 to 0 mM substrate transitions into four groups––depending on the time spent on 500 mM prior to the transition––we found that the average, “baseline” ΔF/F on 500 mM became progressively higher. After > 3 min. on 500 mM, the mean ΔF/F on 500 and 0 mM became indistinguishable (Fig. 4c). It is intriguing that this slow increase in oviDN mean [Ca^2+^] in flies residing on a 500 mM substrate occurs on a minutes timescale that roughly matches the timescale over which egg-laying rates recover in flies residing on 500 mM in free behavior (compare Fig. 4c with Fig. 3f). Consistent with the idea that oviDN [Ca^2+^] tracks relative value and not just sucrose concentration, the magnitude of the average ΔF/F changes during substrate transitions from 0 to 500, from 0 to 200, and from 200 to 500 mM were similar (Extended Data Fig. 13).

We hypothesize that excitatory inputs associated with the relative value of the current substrate interact with additional, more generic, excitatory drive associated with the search state. The generic inputs might start arriving to oviDNs after the search start arrow in Figure 1k, where one observes a small inflection in the oviDN ΔF/F. These two inputs ultimately drive the oviDNs to hit threshold, thus inducing egg laying. One prediction of this model is that the slope at which the oviDN [Ca^2+^] signal approaches threshold should be shallower on the less valued substrate, because of reduced drive from the putative relative-value inputs, and steeper on the more valued substrates. Although the number of eggs available to analyze were very few, we did find, consistent with our hypothesis, that the mean slope of the oviDN ΔF/F rise-to-threshold was shallower on the lower relative-value substrate than the higher one (Fig. 4d) and this was also evident, to near statistical significance, in an analysis of individual traces (Fig. 4e). These data are inconsistent with an alternative hypothesis where the threshold’s height is inversely related to relative value (Fig. 4d), for example. In free behavior, we would expect the slope of the subthreshold oviDN signal to show even more marked adjustments than those shown in Figure 4d, because, unlike head-fixed flies, freely walking flies transition between low and high relative-value substrates regularly during search.

## Gentle hyperpolarization of oviDNs increases the search duration resulting in improved choice performance

Given the above framework for how the oviDN signal guides egg-laying substrate choice (Fig. 4a), we asked whether we could perturb oviDNs to induce flies to be more patient and make “better” choices. Flies lay eggs on the option with lower relative value (higher sucrose) if they have not visited the more valued option in the past ∼2 min. (Fig. 3f-h). We hypothesized that if we could increase the time it takes for [Ca^2+^] to reach threshold by gently hyperpolarizing all ten oviDNs, flies would have more time to encounter the higher relative-value option and may thus lay eggs on the higher value option even more often than normal. Anthropomorphizing, these flies might be considered as more patient, trading off decision speed for increased accuracy.

Expressing the human Kir2.1^34^ potassium channel in oviDNs eliminates egg laying^11^ (Fig. 5a), as does optogenetic inhibition restricted to the egg-laying assay period using the light-gated anion channel, GtACR1^35^ (Fig. 5b, Extended Data Fig. 14). We presume that each of these perturbations prevents the oviDN signal from ever hitting threshold. We found a modified mouse Kir2.1 (hereafter Kir2.1*) that, when expressed in all ten oviDNs, still permits egg laying, albeit at lower average amounts (Fig. 5c). If we only consider flies that laid ≥ 5 eggs, the mean number of eggs per fly becomes more similar, 37 vs. 29, suggesting that Kir2.1* may prevent threshold from ever being reached in some individuals but not others. We used genetic-background matched control flies that expressed a non-conducting channel, Kir2.1*Mut, which differs from Kir2.1* by only 3 amino acids^36^. Kir2.1* and Kir2.1*Mut had been developed and used to hyperpolarize neurons in mice^36^; we introduce the transgenes into flies here (Methods). A similar strategy of using Kir2.1 paired with a non-conducting control has been recently used in flies^37^, but in those experiments the human, not a modified mouse, isoform was used. Kir2.1* and Kir2.1*Mut gene expression was restricted to a 23-hour period prior to experiments (Methods). We performed whole-cell patch-clamp recordings from oviDNs and found that Kir2.1* expressing oviDNs were hyperpolarized by ∼14 mV, on average, compared to Kir2.1*Mut expressing flies (Fig. 5d). This is a moderate hyperpolarization that still permitted most Kir2.1* expressing oviDNs to fire spikes with sufficient current injection (Extended Data Fig. 15a). This fact could explain why at least some Kir2.1* flies can lay eggs.

**Fig. 5.**
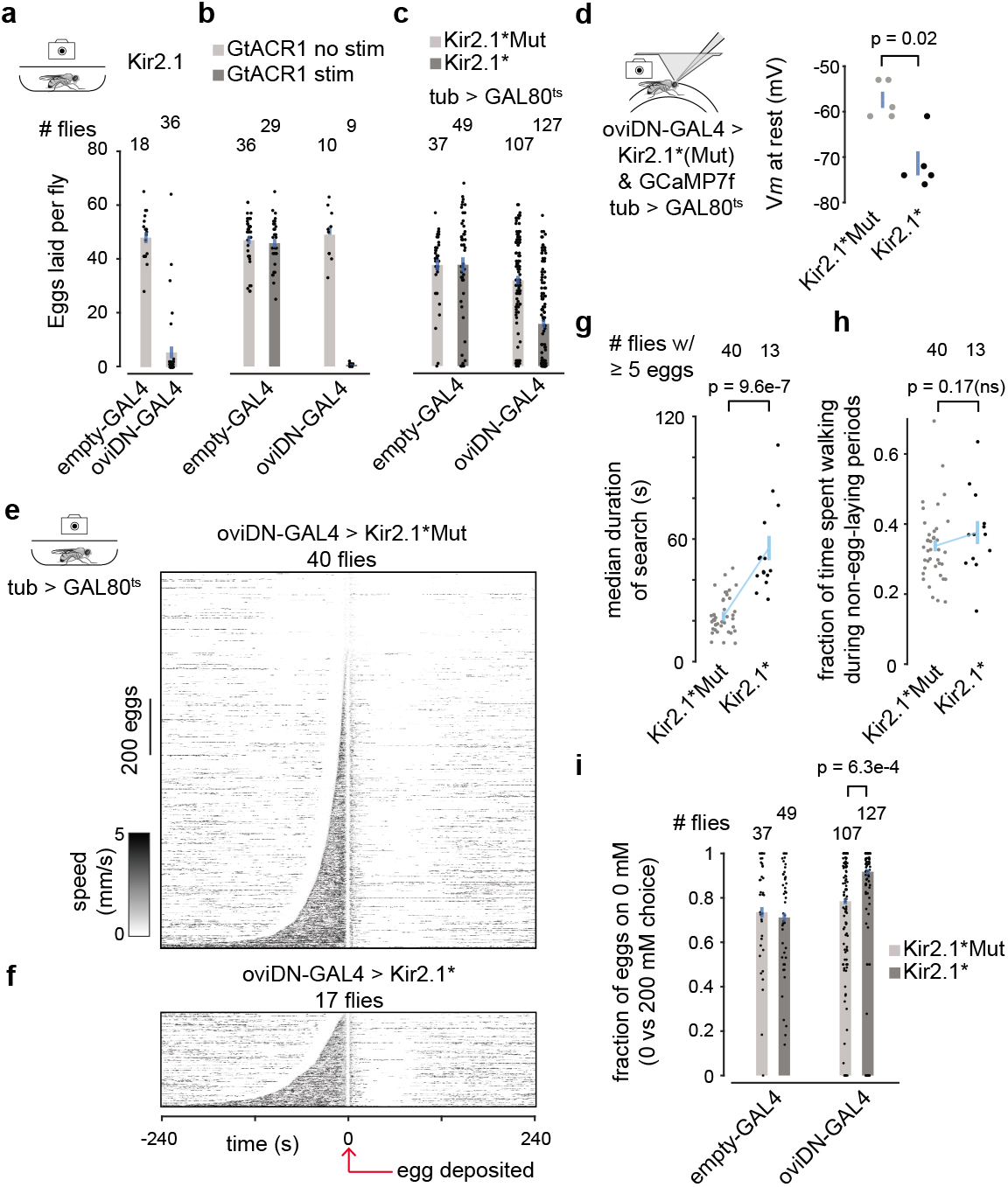
Gentle hyperpolarization of oviDNs increases the search duration and results in more accurate decisions. **a-c,** Eggs laid per fly. Each dot is one fly. **d,** oviDN *Vm* at rest. 5 cells in 5 flies and 5 cells in 4 flies, respectively. **e, f,** Each row represents a single egg-laying event in a 0 vs. 200 mM sucrose chamber, aligned to egg deposition, with the fly’s speed indicated by the black intensity. 1377 eggs from 40 flies (45 flies tested and 5 did not lay eggs) and 346 eggs from 17 flies (40 flies tested and 23 did not lay eggs), respectively. **g,** Median duration of search period for individual flies that laid ≥ 5 eggs from panels e-f. **h,** Fraction of time spent walking during non-egg-laying periods for the same flies as in panel g. Non-egg-laying periods were defined as moments > 10 min from egg deposition. **i,** Fraction of eggs on the lower sucrose option with 95% confidence interval. Each dot is one fly. If the same plot shown is redone by examining only flies that laid ≥ 5 eggs then p = 2.6e-6. In panels c-i, tub > GAL80^ts^ was present in all flies to limit the time window over which Kir2.1* or Kir2.1*Mut transgenes were expressed (Methods).

We tracked the x-y trajectories and egg-laying behavior of flies expressing Kir2.1* or Kir2.1*Mut in all ten oviDNs (using the oviDN-GAL4) in two-substrate-choice, free-behavior chambers. We observed a 2-3 fold increase in the search duration in Kir2.1* compared to Kir2.1*Mut flies when comparing the distribution of individual traces by eye (Fig. 5e-f) or when quantifying the median search duration (Fig. 5g). The increase in search duration cannot be attributed to a general increase in the fraction of time spent walking (Fig. 5h) or a broad defect in egg-laying-related motor functions (Extended Data Fig. 15b-c). An increase in search duration when all oviDNs are gently hyperpolarized is presumably related to [Ca^2+^] taking longer to reach threshold in oviDNs. This increase in search duration was, remarkably, accompanied by a higher fraction of eggs laid on the substrate of higher relative value, i.e., an improvement in choice behavior (Fig. 5i). Note that flies expressing Kir2.1*Mut in oviDNs expressed a weaker baseline trend for laying eggs on low sucrose (Fig. 5i, right light grey bar) compared to Canton S flies (Fig. 3b); however, ∼75-80% choice is comparable to the level exhibited by our empty-Gal4 controls (Fig. 5i, left light grey bar) and is a common preference level for many genotypes (data not shown from a second manuscript *in preparation*).

## Discussion

We provide insight into the cellular mechanisms by which animals perform ethologically relevant, relative-value-based decisions. Specifically, dynamic modulation of oviDN [Ca^2+^] over the course of many seconds to minutes determines when that neuron’s activity hits a threshold and we propose that this process fundamentally regulates where flies lay eggs.

Our work describes a rise-to-threshold [Ca^2+^] signal in the oviDN soma. Although this soma [Ca^2+^] signal could have an important function (e.g., controlling a transcriptional process^38^ that keeps the mean oviDN [Ca^2+^] levels within a specific range), it itself cannot drive motor action because the soma is far from the output synapses of the neuron. In unipolar neurons, like oviDNs, changes in soma [Ca^2+^] are typically caused by *Vm* fluctuations elsewhere in the neuron propagating back to the cell body and inducing voltage-gated calcium channels to open^39^. Thus, the presumed *Vm* fluctuations that caused our soma [Ca^2+^] signal to change are also expected to propagate down to the oviDN synapses in the abdominal ganglion. Voltage-gated calcium channels at the presynaptic terminals in the abdominal ganglion are also likely to open, potentially creating a rising [Ca^2+^] signal in the terminals that resembles the one we describe in the soma. Moreover, given the sharp, non-linearity between presynaptic [Ca^2+^] and synaptic vesicle release^40^, one can imagine how a threshold might actually be implemented in presynaptic terminals. Alternatively, the oviDNs may transmit a graded signal to post-synaptic partners resulting in those neurons implementing the threshold. Future work will be needed to link the rise-to-threshold signal evident in the oviDN soma, mechanistically, to motor action.

A fly’s decision of when and where to lay an egg resembles decisions that humans make, like when to stop browsing a menu and choose a dish at a restaurant. Both processes start with an initiation event: ovulation or the decision to have a meal. Next, the individual’s own behavior reveals new options to the organism over time: more egg-laying substrates or more dishes. Finally, the decision process is terminated when one option is selected and the motor program to choose that option has begun. This structure would seem to apply broadly, to many natural decisions taken by animals and humans, and thus the principles by which the oviDN mediates *Drosophila* egg laying may prove relevant for understanding decision making more broadly, across contexts and species.

Rise-to-threshold signals have been linked to decision-making and action initiation in humans^41, 42^, monkeys^4, 43–46^, rodents^47–52^, zebrafish^53–55^, and insects^56–60^. These signals have been shown to rise, or hypothesized to rise, on the hundreds-of-milliseconds to seconds timescale. In the decision-making literature, some of the most influential work has focused on so-called “perceptual decisions”, in which rising signals integrate sensory input so that an animal can report a percept^2, 3^. More recently, neural signals associated with self-paced decisions have been studied^5–8^, but without definitive links to behavior yet. The oviDN signal we describe emphasizes several important features of how brains make decisions. (1) Rise-to-threshold signals can underlie decisions that take minutes, not just seconds. (2) Rise-to-threshold signals can cause behavior to start when threshold is crossed^50, 59^; they are not just a correlate of the decision-making process. (3) Rise-to-threshold signals can track the relative value of stimuli to guide their ramping rate, not just the sensory properties of those stimuli. (4) Rise-to-threshold signals underlie ethologically relevant, self-paced decisions––where options are encountered due to the animal’s own actions––not just externally cued decisions. This latter finding is particularly important because it generalizes the rise-to-threshold mechanism to the vast swath of decisions that animals make where their own actions determine the options experienced.

## Extended Data, Supplemental Materials, and Methods

### Extended Data Legends

**Extended Data Fig. 1.**
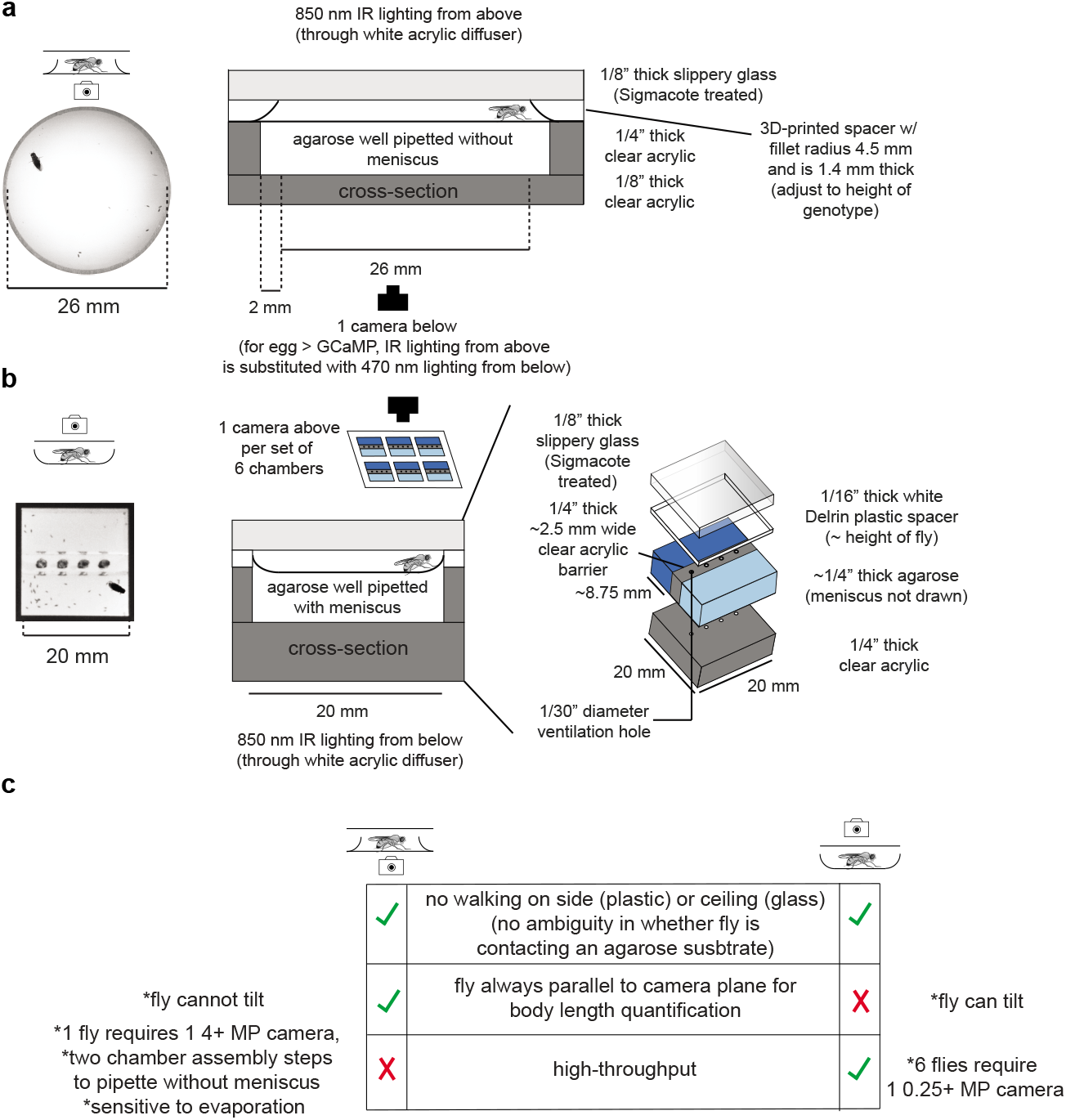
Free behavior chambers. **a,** Schematic of free behavior egg-laying chamber with sloped ceiling. **b,** Schematic of high-throughput free behavior egg-laying choice chamber. **c,** Comparison of the two free behavior chamber types.

**Extended Data Fig. 2.**
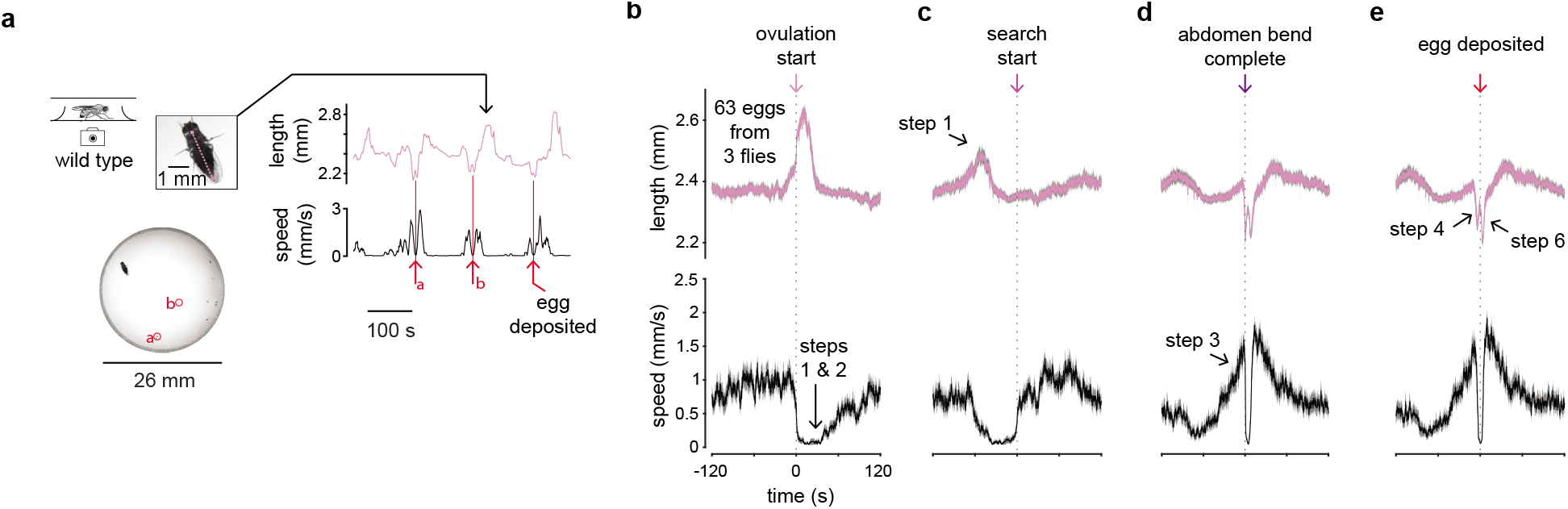
Characterization of the egg-laying behavioral sequence. **a,** Length (neck to ovipositor distance) and locomotor speed over 3 consecutive egg-laying events, smoothed with a 5 s boxcar filter, for a single fly in a sloped ceiling egg-laying chamber (Supplementary Video 2). **b-e,** Length and locomotor speed aligned to annotated events in the egg-laying behavioral sequence. Prominent features of steps from Fig. 1a that were not considered when annotating the event used for alignment are labeled.

**Extended Data Fig. 3.**
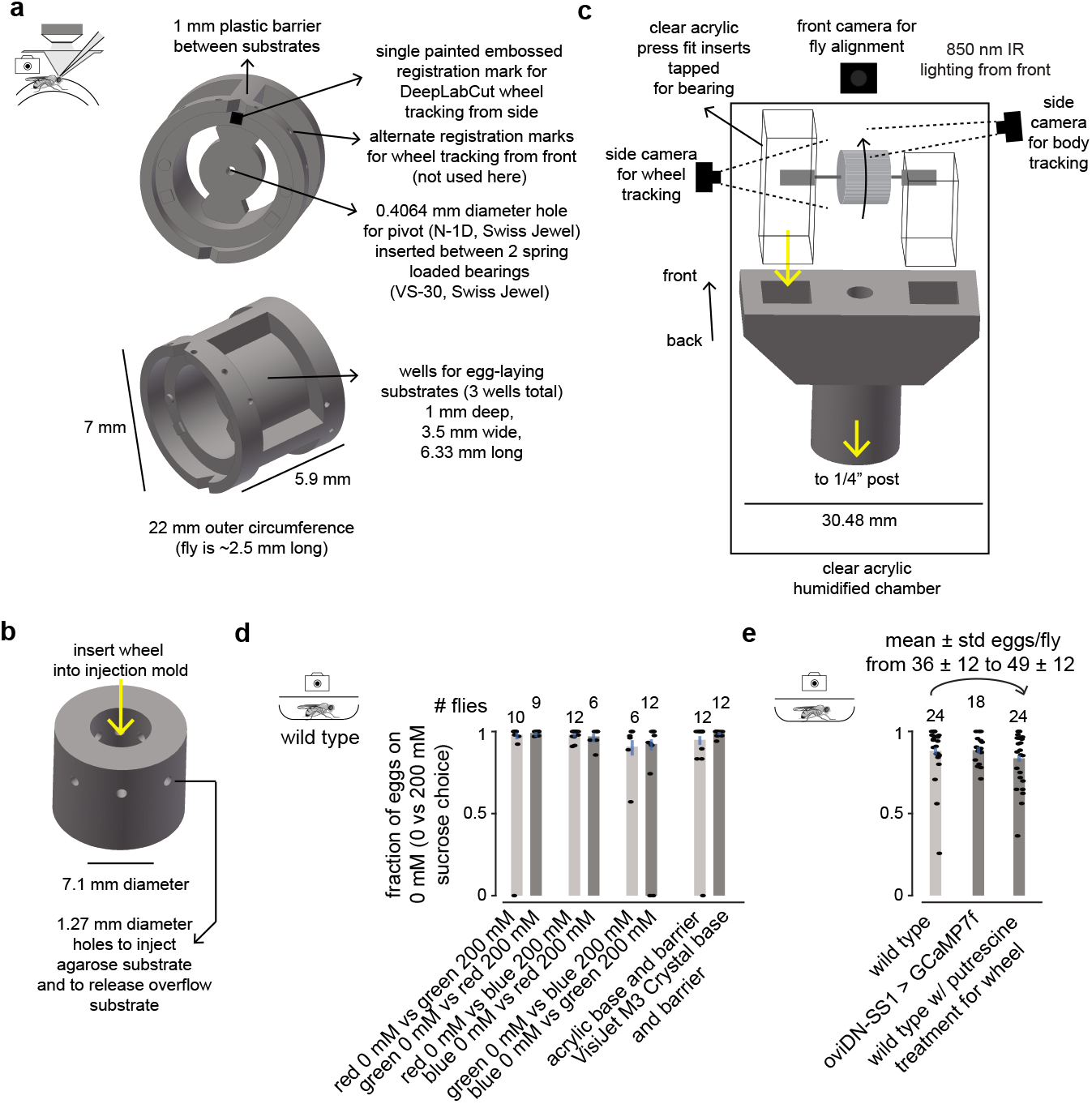
Egg-laying wheel design and behavior in control egg-laying chambers and by control flies. **a,** Schematic of egg-laying wheel. **b,** Schematic of agarose-injecting mold, which is used to load agarose onto the wheel using a pipette. **c,** Schematic of egg-laying wheel assembly secured in a custom humidification chamber under the microscope objective. **d,** Fraction of eggs on the lower sucrose option for control substrates: colored dye infused substrates and 3D-printer material (VisiJet M3 Crystal) bases vs. acrylic bases. Error bars represent 95% confidence intervals. Each dot is one fly. **e,** Fraction of eggs on the lower sucrose option for flies expressing GCaMP7f in oviDNs and by those pre-treated for tethered wheel experiments. Error bars represent 95% confidence intervals. Each dot is one fly.

**Extended Data Fig. 4.**
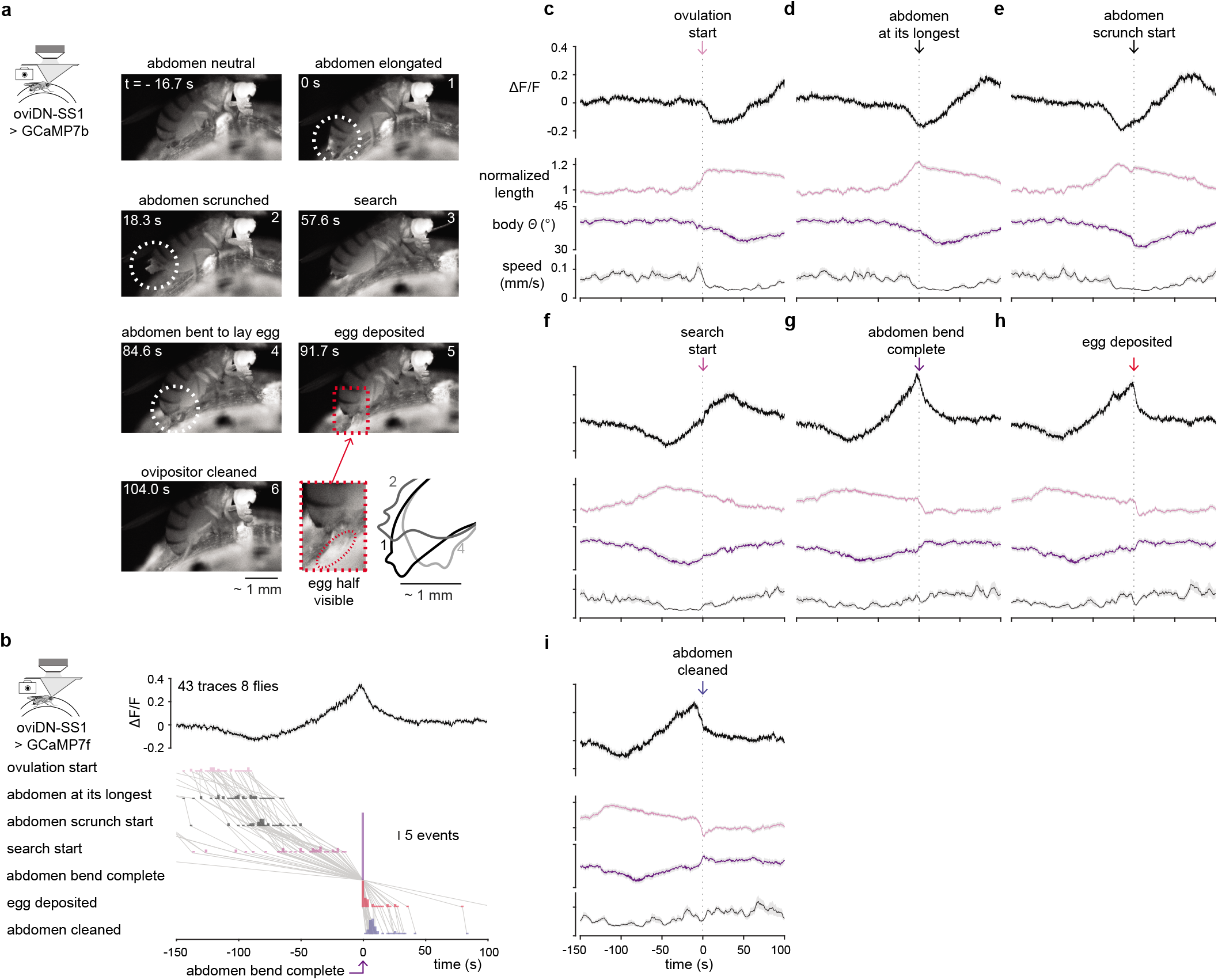
Tethered egg-laying behavioral sequence with oviDN [Ca^2+^]. **a,** Behavioral sequence of tethered egg laying as in Figure 1a. Stills from a single egg-laying event. Overlaid and zoomed-in schematics of the tip of the abdomen from 3 frames is shown at the bottom right. **b,** Mean oviDN ΔF/F aligned to the moment abdomen bending to lay an egg is complete. 43 traces from 9 cells in 8 flies (41 eggs). Behavior shown below. **c-i,** Mean oviDN ΔF/F and behavior aligned to events in the behavioral sequence shown in panel b. Locomotor speed is smoothed with a 5 s boxcar filter.

**Extended Data Fig. 5.**
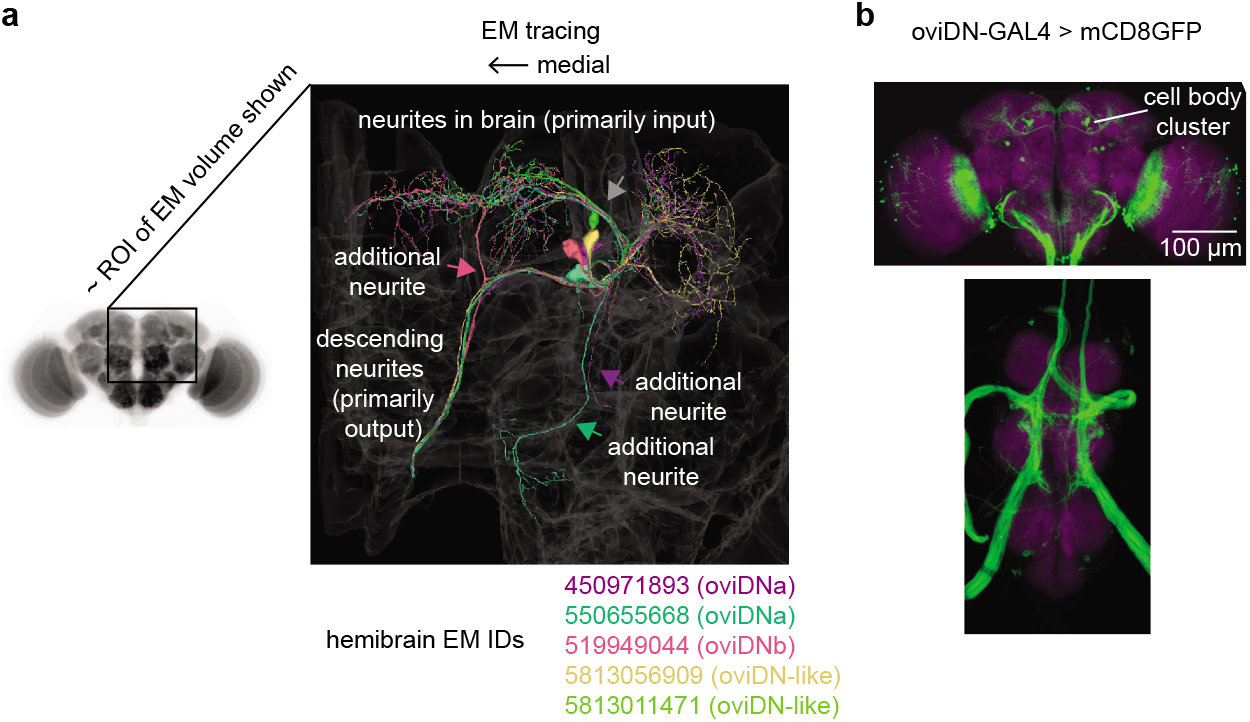
Anatomy of oviDNs. **a,** Electron-microscopy (EM) tracing^29^ and characterization of the 3 oviDN and 2 oviDN-like neurons per side. The branch labeled in grey is sometimes present in oviDNb^11^ and sometimes not (Fig. 1e). The 3 other arrows indicate neurites that are unique to either oviDNa or oviDNb. Visualization generated using Neuroglancer. Neuropil to left is only to schematize the approximate ROI shown in the EM. **b,** Average z-projection of oviDN-GAL4 in the brain (top) and ventral nerve cord (bottom).

**Extended Data Fig. 6.**
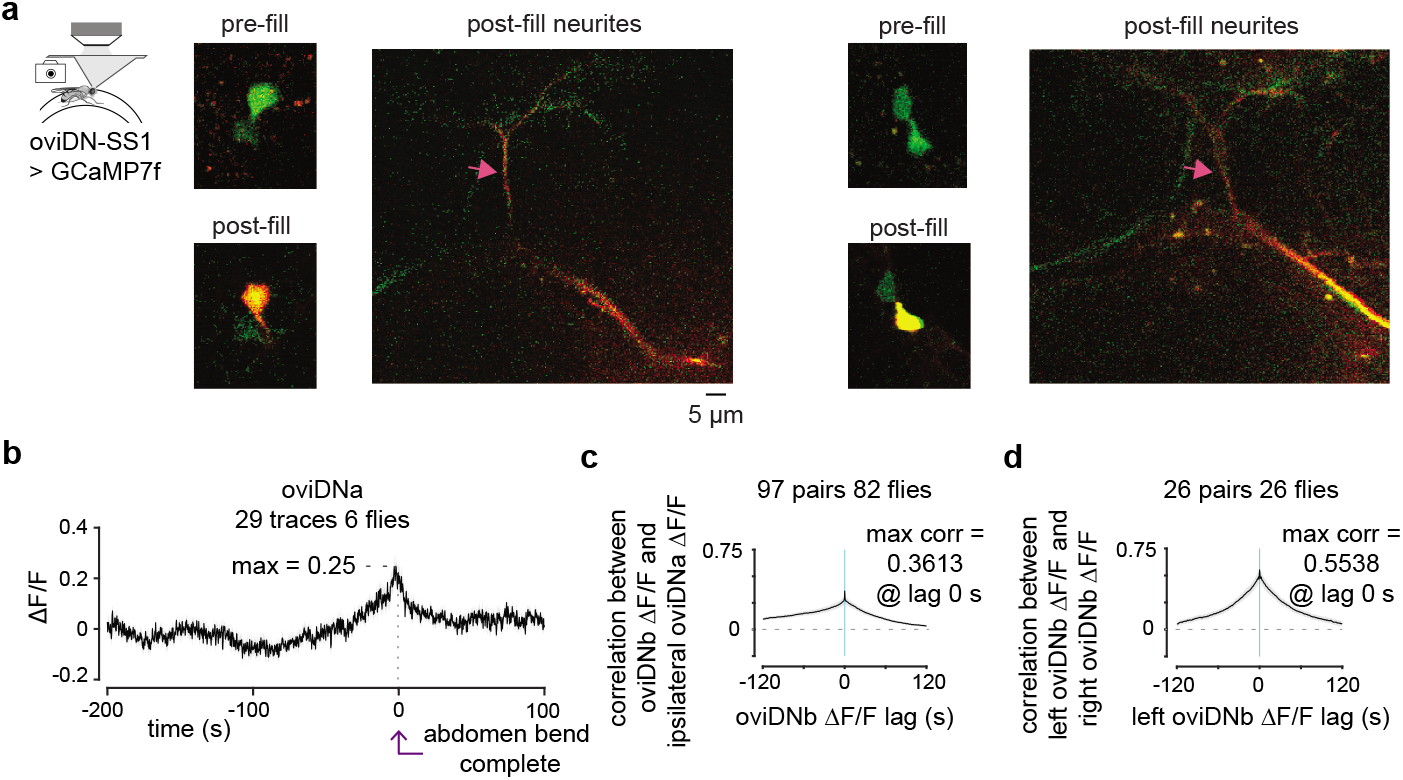
oviDNa and oviDNb have qualitatively similar physiology. **a,** Anatomy of oviDN-SS1 driving expression of GCaMP7f. The brighter of the two oviDN cell bodies was filled with Texas Red (Methods). The neurite labeled with a pink arrow (see Extended Data Fig. 5a) was used to determine if the cell was oviDNb. All 6 of the brighter cells filled with Texas Red (from 6 separate flies) were oviDNb. Two examples are shown (representative individual z-slices). **b,** ean oviDNa ΔF/F during individual egg-laying events. 29 traces from 7 cells in 6 flies (28 eggs). **c,** Mean correlation of ΔF/F between ipsilateral oviDNa and oviDNb cells imaged simultaneously. Individual cell pairs are averaged. **d,** Mean correlation of ΔF/F between contralateral oviDNb cells imaged simultaneously. Individual cell pairs are averaged.

**Extended Data Fig. 7.**
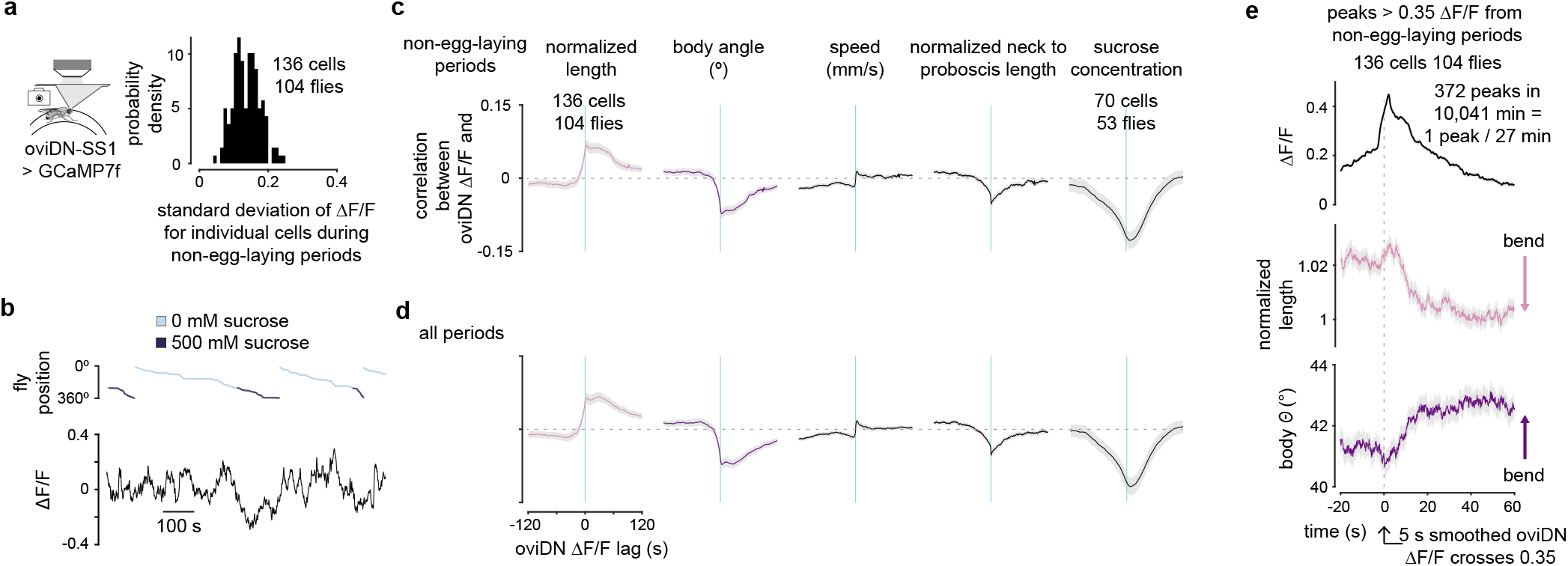
oviDN ΔF/F fluctuates during non-egg-laying periods and these fluctuations are correlated to behavior. **a,** Standard deviation of oviDN ΔF/F during non-egg-laying periods. Non-egg-laying periods were defined as moments > 5 min away from egg deposition. **b,** OviDN ΔF/F during a non-egg-laying period (smoothed with a 2 s boxcar filter) and behavior. This cell had a standard deviation in ΔF/F of 0.15. **c,** Mean correlation of oviDN ΔF/F and behavior during non-egg-laying periods. For sucrose concentration correlations, only 0 vs. 500 mM sucrose wheels were analyzed (excluding 0 mM only wheels, for example), leaving only 53/104 flies for analysis. **d,** Same as panel c, but including time periods near egg deposition. **e,** Mean oviDN ΔF/F and behavior during peaks in ΔF/F that occurred in non-egg-laying periods. We smoothed the ΔF/F signal with a 5 s boxcar filter and extracted peaks in the ΔF/F trace that exceeded 0.35 for > 1 s. We aligned these traces to the moment the ΔF/F signal crossed 0.35 in the 10 s prior to the peak.

**Extended Data Fig. 8.**
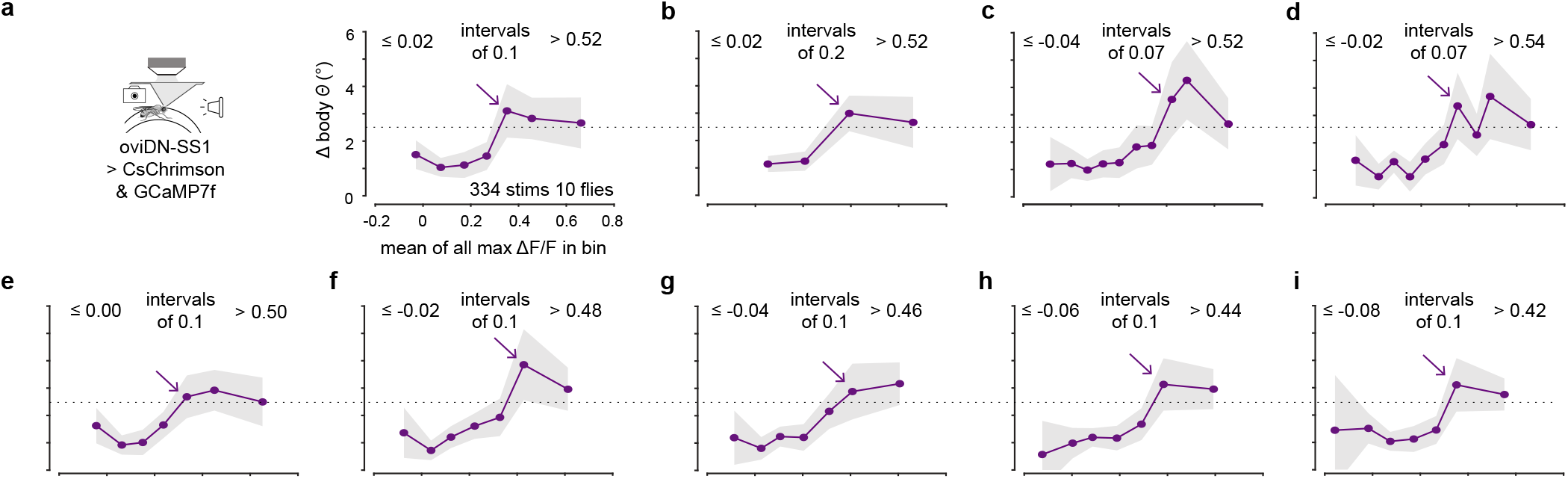
Varied binning for abdomen-bend vs. oviDN ΔF/F support the hypothesis that a threshold in oviDN [Ca^2+^] activity triggers the egg-deposition motor program. **a,** Change in mean body angle, replotted from Figure 2h. Arrow indicates first bin with an abdomen angle change greater than 2.5° (indicated by dotted line). **b,** Same as panel a but with coarser binning. **c, d,** Same as panel a but with finer binning. **e-i,** Same as panel a but bins are shifted progressively by 0.02 leftward. **a-i,** The first and last bin always include all the data points below and above that bin, respectively. The curve in panel g appears more linear and less step-like than the others; however, it is expected that as one progressively shifts the center point of the bins, one will find a position where the central bin exactly straddles the putative threshold, yielding an intermediate y value for that bin. The fact that panels f and h appear more step like supports this explanation for panel g.

**Extended Data Fig. 9.**
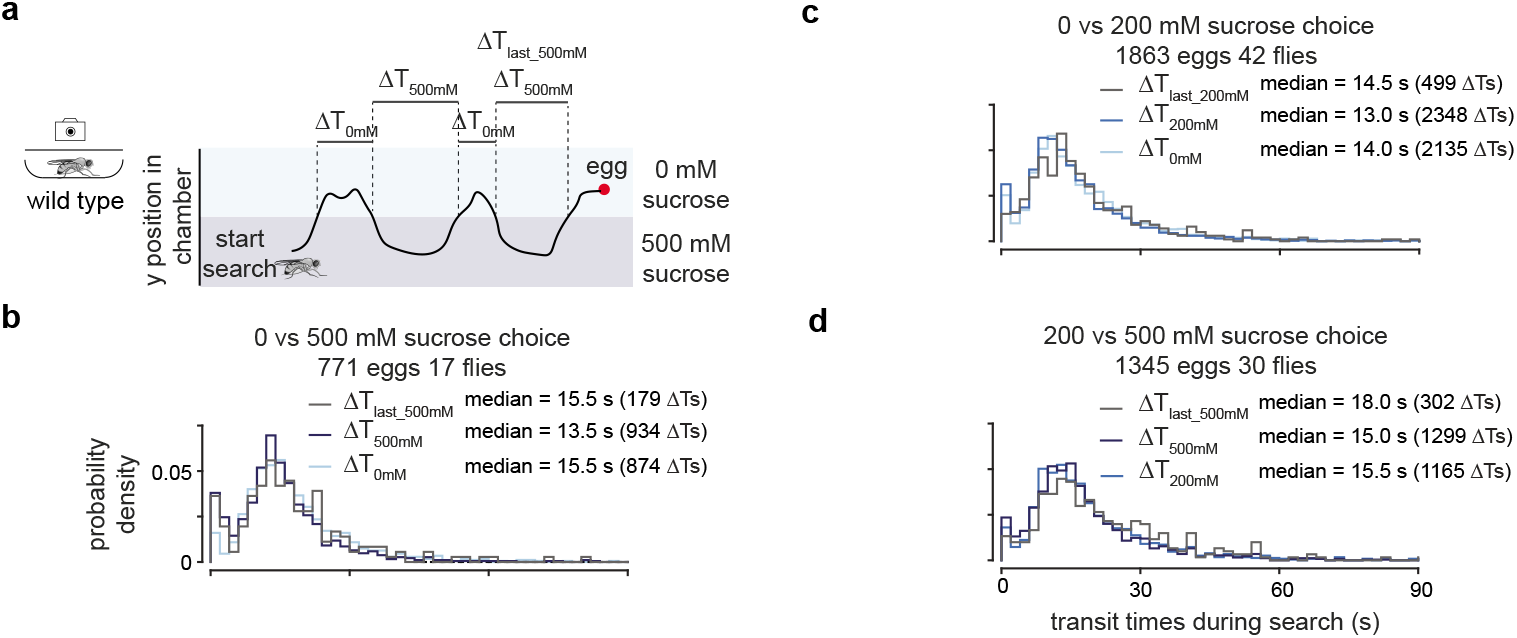
Evidence against use of spatial information guiding search and against feeding-on-higher-sucrose-related suppression of egg deposition in our free behavior chambers. **a,** Schematic of a fly searching for an egg deposition site in a 0 vs. 500 mM chamber. ΔT _0mM_ and ΔT_500mM_ are all the intervals of time that a fly was transiting through 0 or 500 mM, respectively, during an egg-laying search period. ΔT_last_500mM_ is the last transit interval through 500 mM for eggs deposited on 0 mM. If a fly were positionally avoiding sucrose, ΔT_500mM_ would be less than ΔT_0mM_. If a fly were to use spatial information during the search period––by taking a shortcut to get to the preferred 0 mM substrate at the end of a search––ΔT_last_500mM_ would be less than ΔT_0mM_ and ΔT_500mM_. If a fly were feeding on the higher sucrose substrate – and pausing as flies do when they feed^61^––ΔT_500mM_ would be larger than ΔT_0mM_. **b-d,** ΔT_lower_sucrose_, ΔT_higher_sucrose_, and ΔT_last_higher_sucrose_ distributions for three different sucrose choice chambers. ΔT_higher_sucrose_ is not less than ΔT_lower_sucrose_ suggesting that flies are not positionally avoiding the higher sucrose option. ΔT_last_higher_sucrose_ is not detectably smaller than ΔT_0mM_ or ΔT_500mM_ suggesting that flies are not taking a shortcut––and thus not manifesting use of spatial information––at the end of the search. It is possible that flies use spatial information to guide the search in conditions with less thigmotactic walking^62, 63^ and/or with visual landmarks^64^ (all experiments in this study were conducted in darkness). Note that use of spatial information is compatible with our time-domain model for egg laying and would just suggest that flies control the substrate upon which they are located and as such control the egg-laying drive that they experience. ΔT_higher_sucrose_ is not larger than ΔT_lower_sucrose_ indicating that flies are not pausing only on the higher sucrose substrate. We interpret this result to mean that flies are not suppressing egg deposition because of extensive feeding on the sucrose substrates. In addition, we did not notice additional proboscis extension on higher sucrose when we spent hours inspecting each egg to annotate the egg deposition time, probably because our flies are very well fed prior to entering the chamber (Methods). 771 eggs from 17 flies (18 flies tested and 1 did not lay eggs), 1863 eggs from 42 flies (47 flies tested and 5 did not lay eggs), and 1345 eggs from 30 flies (30 flies tested), respectively.

**Extended Data Fig. 10.**
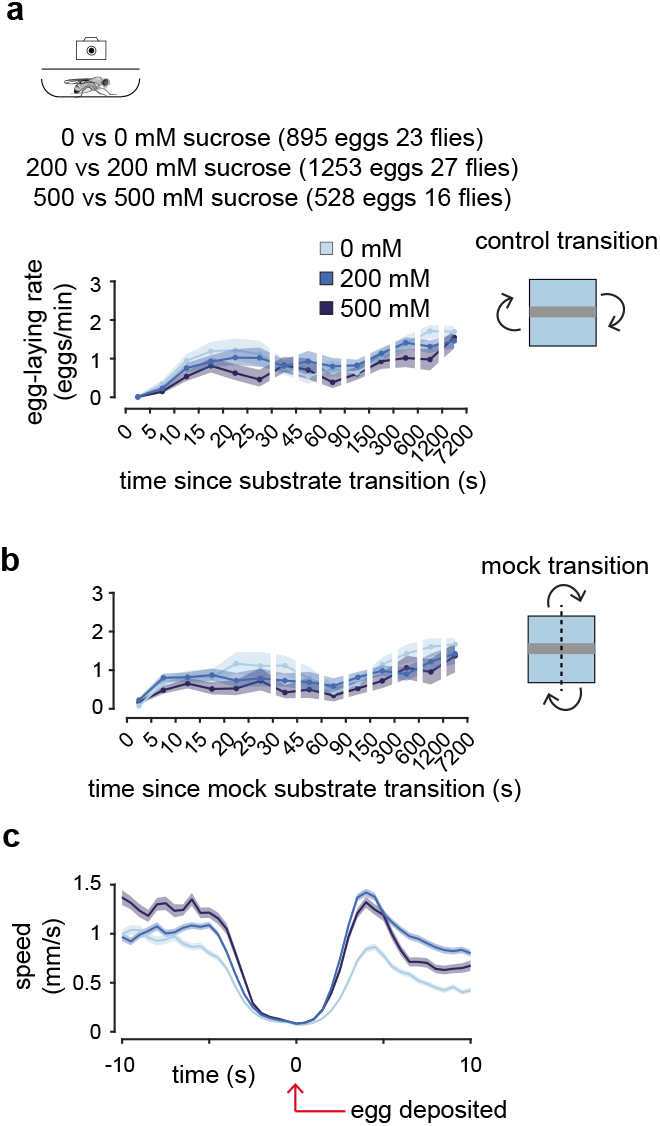
Controls for egg-laying rate function. **a,** Mean egg-laying rates during the search period after a fly transitions across the plastic barrier in a single-option chamber, meaning that there is either 0 mM sucrose on both sides, 200 mM sucrose on both sides, or 500 mM sucrose on both sides. 90% confidence interval shaded. Egg-laying rates on the three different sucrose concentrations are similar in single-option chambers. A slight trend for higher egg-laying rates on lower sucrose suggests a slight innate preference for lower sucrose may exist. This slight innate preference cannot explain the much larger differences in rate in two choice chambers (Fig. 3f-h). 895 eggs from 23 flies (24 flies tested and 1 laid no eggs), 1253 eggs from 27 flies (27 flies tested), and 528 eggs from 16 flies (17 flies tested and 1 laid no eggs) for 0 vs. 0, 200 vs. 200, and 500 vs. 500 mM chambers, respectively. **b,** Mean egg-laying rate during the search after a fly transitions across a mock vertical line. Same data as in panel a. The 5-10 s bin in this analysis has a higher egg laying rate than in the analysis from panel a, suggesting that part of the delay in egg laying after a transition is due to flies not laying eggs on the plastic barrier. **c,** Mean locomotor speed. A ∼3 s delay exists between when a fly pauses and bends its abdomen to lay an egg till when an egg is deposited. This ∼3 s can explain why even the data in panel b do not show high egg laying rates in the 0-5 s bin. Analyzing the same data as in panels a-b.

**Extended Data Fig. 11.**
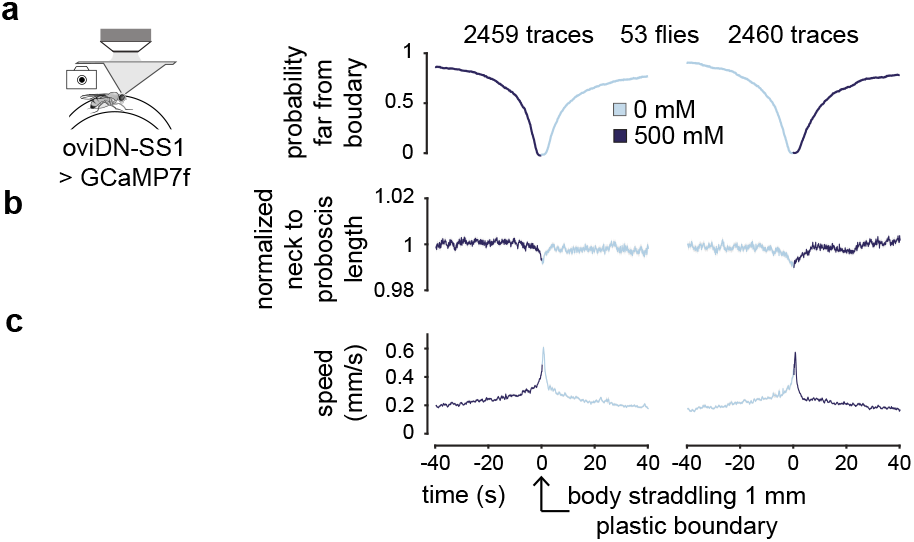
Changes in oviDN ΔF/F during substrate transitions are not due to consistent, detectable changes in behavior. **a,** Probability that a fly’s centroid is > 2 mm from the midline between substrates, aligned to detecting a transition from one substrate to the other. For a 2.5 mm fly this would correspond to the front or back of the fly being 0.75 mm from the midline of the 1 mm plastic barrier between substrates. These data show that it takes a fly ∼10-20 s to completely cross the midline which impacts dynamics measured in our neural signals. 2459 and 2460 traces from 70 cells in 53 flies (1911 and 1922 transitions). **b,** Mean neck to proboscis length during substrate transitions. **c,** Mean locomotor speed during substrate transitions.

**Extended Data Fig. 12.**
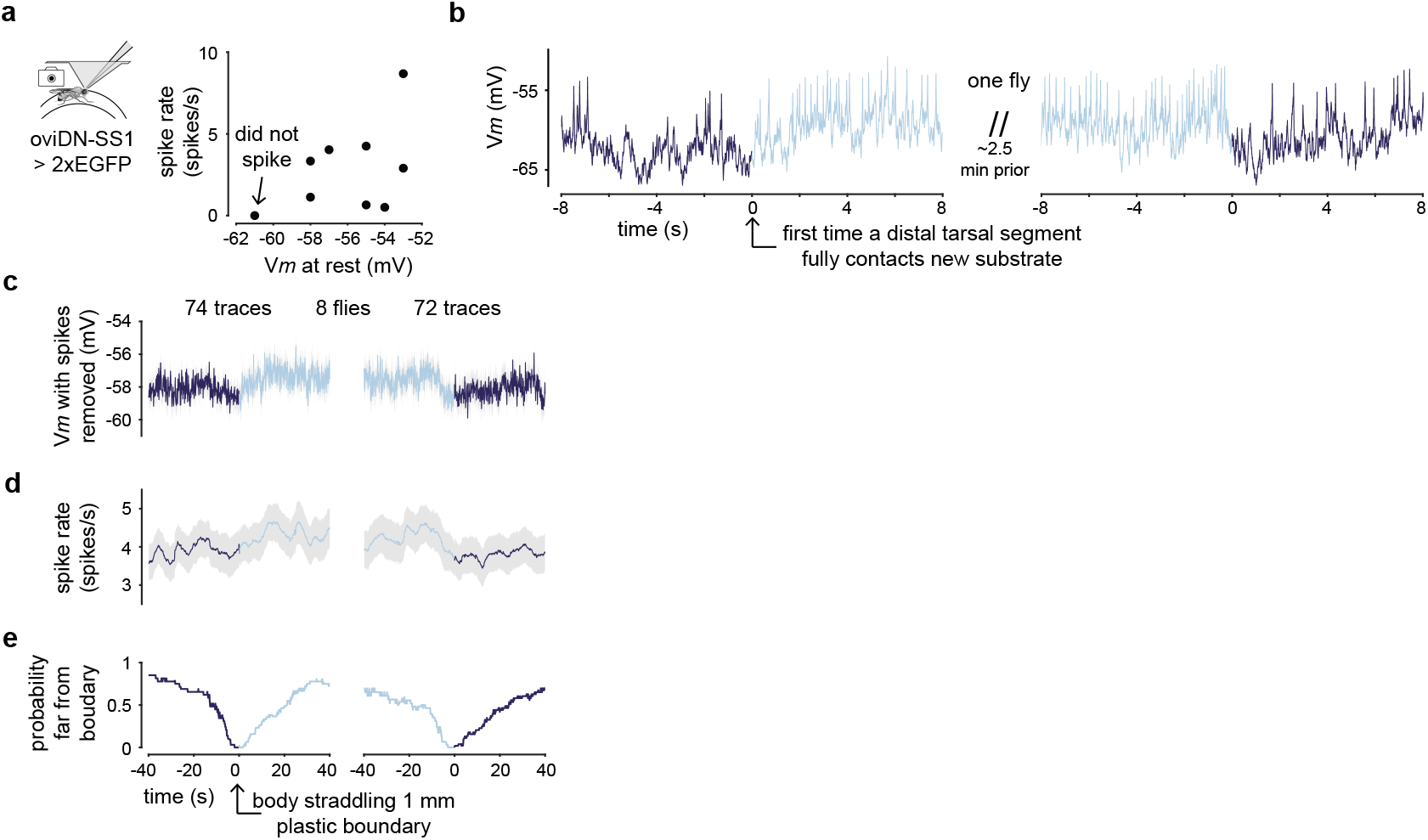
Electrical activity of oviDNs. **a,** oviDN spike rate versus *Vm* at rest. **b,***Vm* during two substrate transitions from the same fly. These sample traces have more pronounced *Vm* changes than is typical. **c,** Mean oviDN *Vm* after removal of spikes during substrate transitions. 74 and 72 traces from 8 cells in 8 flies (74 and 72 transitions). **d,** Mean oviDN spike rate during substrate transitions. **e,** Same as Extended Data Fig. 11a but for electrophysiology dataset.

**Extended Data Fig. 13.**
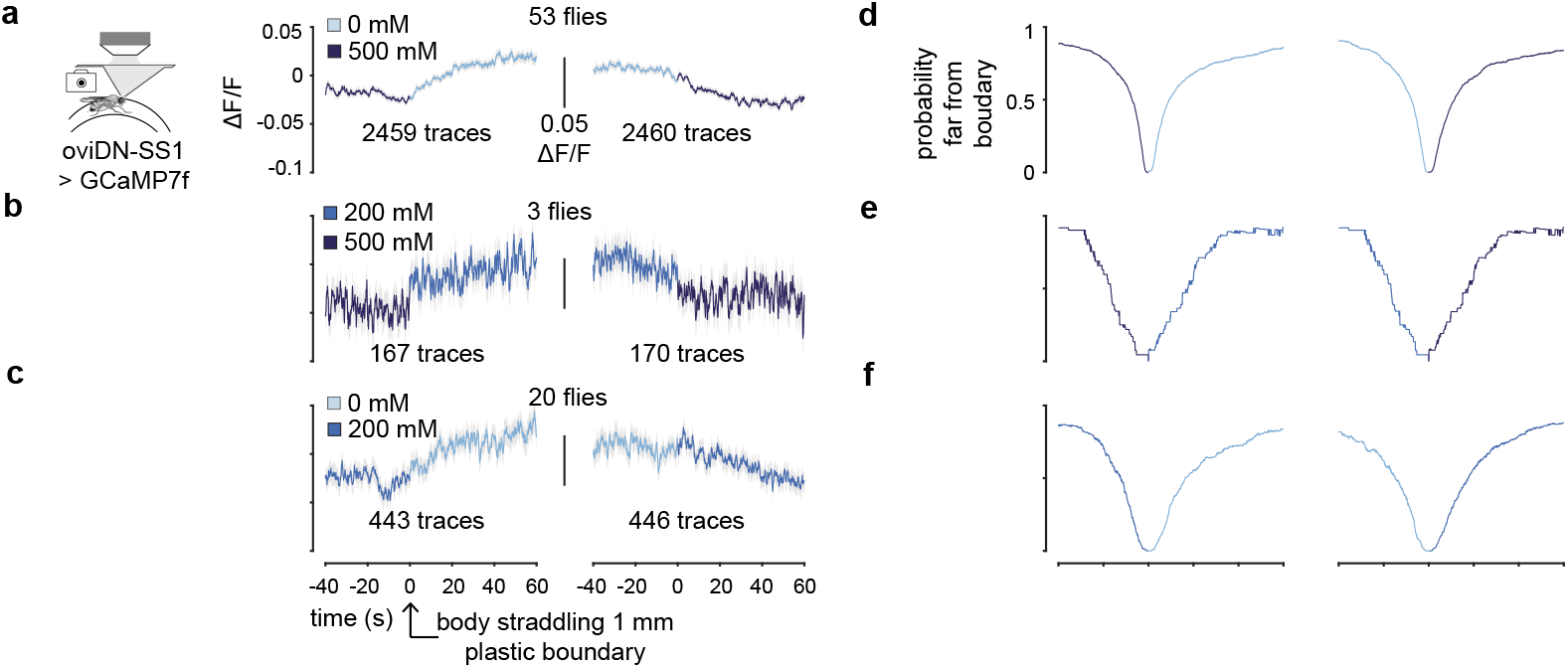
Changes in oviDN ΔF/F during substrate transitions are more consistent with changes tracking relative value than absolute sucrose concentration. **a,** Mean oviDN ΔF/F during substrate transitions from 500 to 0 mM and 0 to 500 mM. 2459 and 2460 traces from 70 cells in 53 flies (1911 and 1922 transitions). **b,** ean oviDN ΔF/F during substrate transitions from 500 to 200 mM and 200 to 500 mM. 167 and 170 traces from 5 cells in 3 flies (105 and 109 transitions). **c,** Mean oviDN ΔF/F during substrate transitions from 200 to 0 mM and 0 to 200 mM. 443 and 446 traces from 20 cells in 20 flies (443 and 446 transitions). In panels a-c, note that all changes are on the order of 0.05 ΔF/F regardless of the absolute sucrose concentration, consistent with a relative value calculation. **d-f,** Same as Extended Data Fig. 11a but for datasets shown to the left.

**Extended Data Fig. 14.**
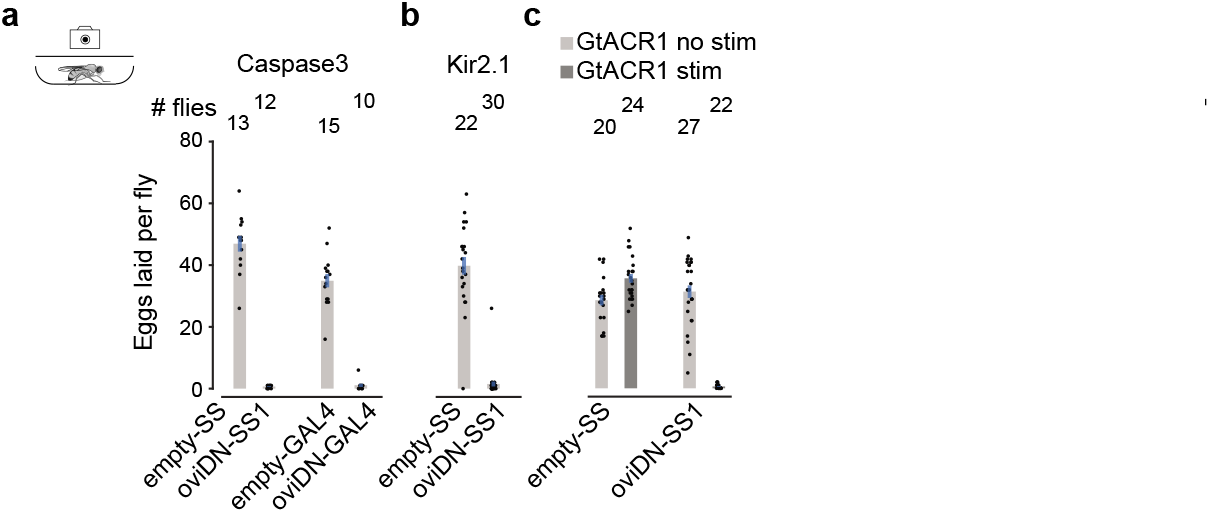
Apoptosis-based, human Kir2.1-based, and optogenetic-based inhibition of oviDN signaling strongly suppress egg deposition. a-c, Eggs laid per fly. Each dot is one fly. 95% confidence interval indicated.

**Extended Data Fig. 15.**
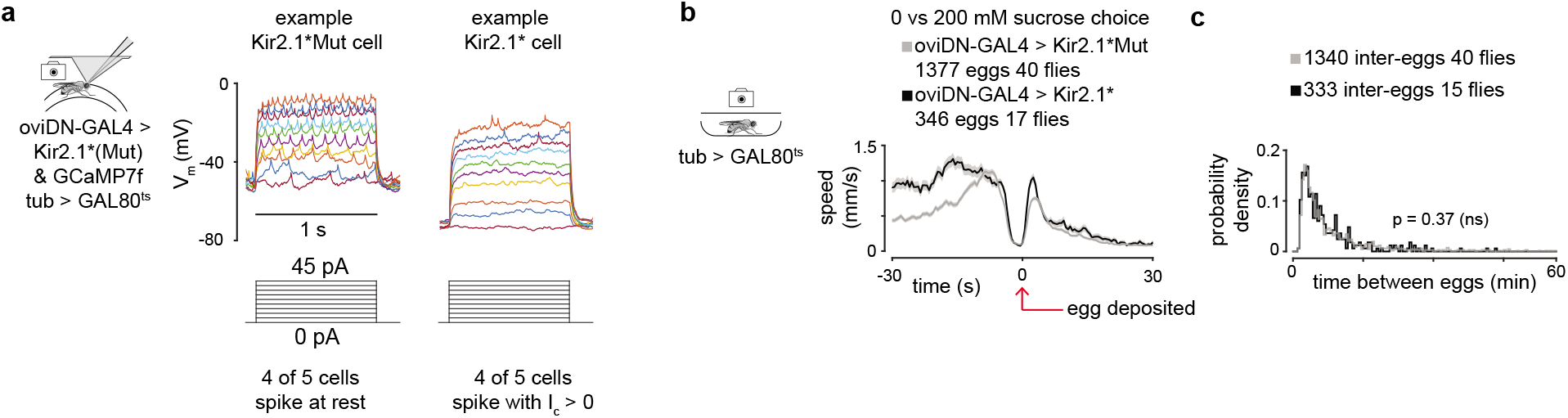
Expression of Kir2.1* in oviDNs does not abolish the ability to spike in most oviDNs and does not significantly alter non-search-related dynamics of egg laying in flies that still lay eggs. **a,** The *Vm* of a single, representative oviDN expressing Kir2.1*Mut or Kir2.1* during current injection. Four out of five Kir2.1* expressing cells showed spikes with sufficient amounts of current injection; one cell did not (not shown). **b,** Mean locomotor speed aligned to egg deposition. A higher average speed prior to egg laying in Kir2.1* flies is indicative of the longer search duration in these flies. However, other aspects like the pause to lay an egg and post-egg-laying speed remain similar in Kir2.1*Mut and Kir2.1* flies. 1377 eggs from 40 flies (45 flies tested and 5 laid no eggs), 346 eggs from 17 flies (40 flies tested and 23 laid no eggs) for Kir2.1*Mut and Kir2.1*, respectively. **c,** Probability density of time between egg-deposition events (inter-egg-intervals). 1340 inter-eggs from 40 flies (45 flies tested and 5 laid < 2 eggs and thus did not have at least 1 inter-egg interval), 333 inter-eggs from 15 flies (40 flies tested and 25 flies laid < 2 eggs and thus did not have at least 1 inter-egg interval) for Kir2.1*Mut and Kir2.1*, respectively. Note that a similar inter-egg interval distribution for Kir2.1* flies and controls does not mean that Kir2.1* flies searched for the same amount of time for an egg-laying substrate as controls; on the contrary, Kir2.1* flies searched longer than controls (Fig. 5g) for a good substrate; it’s just that despite the longer search, they maintained a similar time interval between ovulations. That is, the average time between ovulations, as estimated with locomotor speed, was similar in Kir2.1* and control flies (p = 0.36).

### Supplemental Tables

**Supplementary Table 1.**
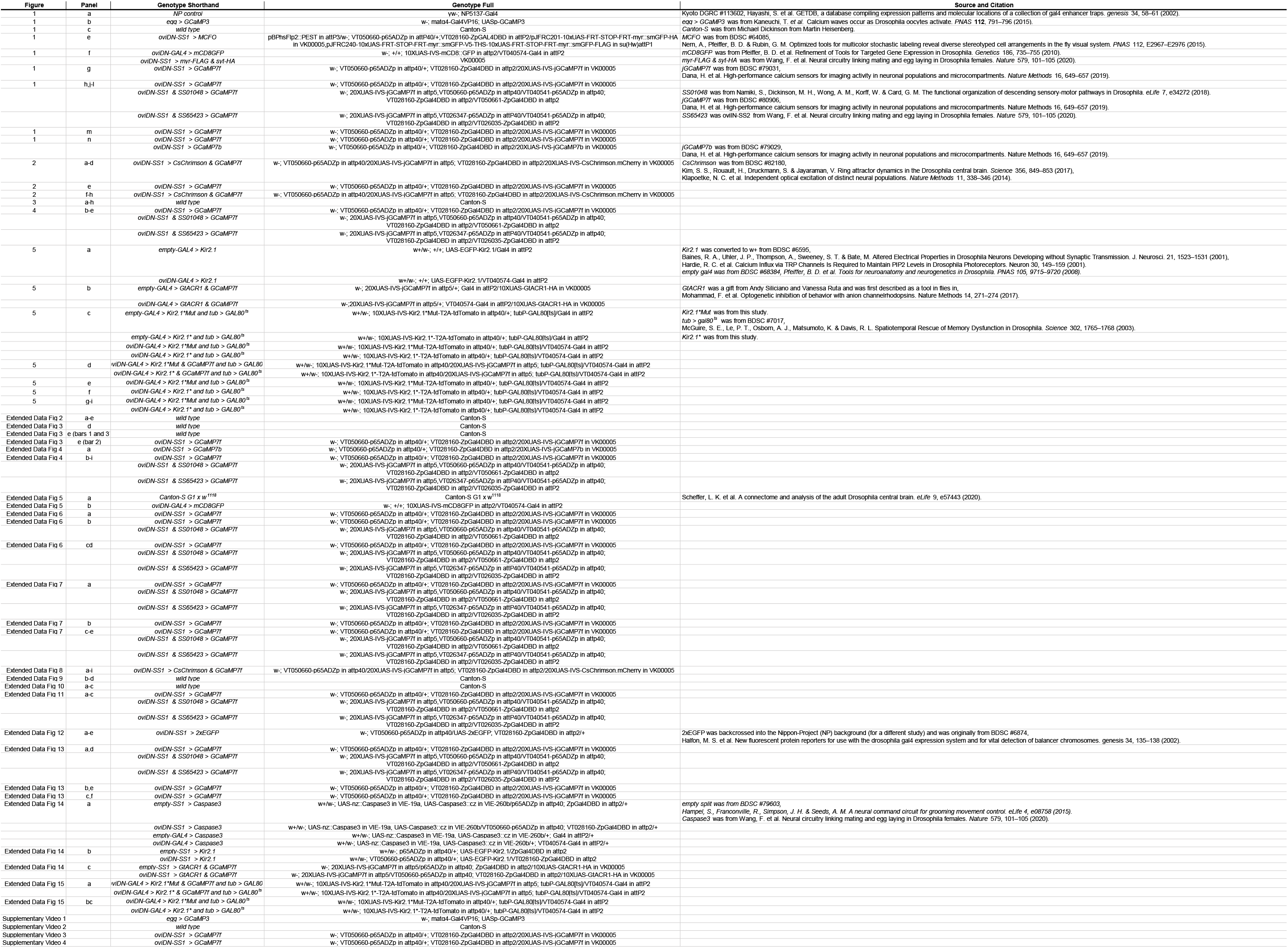
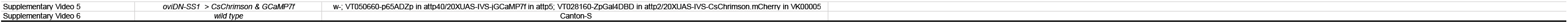
Genotypes for each experiment.

**Supplementary Table 2.** Conditions for each experiment.

| Figure | Panel | Chamber or Wheel | Conditions |
| --- | --- | --- | --- |
| 1 | a | free behavior box | grape juice infused bacto-agar pad |
| 1 | bc | free behavior sloped ceiling | 1% agarose, 0.8% acetic acid, 1.6% ethanol (hereafter, 1% agarose mixture), 0 mM sucrose |
| 1 | g | wheel | 1% agarose mixture, 0 vs 0 vs 500 mM sucrose |
| 1 | h,j-l | wheel | 1% agarose mixture, 0 vs 0 vs 500 mM sucrose<br>1% agarose mixture, 0 vs 500 vs 500 mM sucrose<br>1% agarose mixture, 0 vs 200 vs 500 mM sucrose<br>1% agarose mixture, 0 vs 200 vs 200 mM sucrose<br>1% agarose mixture vs 2% agarose mixture vs 2% agarose mixture |
| 1 | m | wheel | 1% agarose mixture, 0 vs 0 vs 500 mM sucrose |
| 1 | n | wheel | 1% agarose mixture, 0 vs 0 vs 500 mM sucrose<br>1% agarose mixture, 0 vs 200 vs 500 mM sucrose<br>1% agarose mixture, 0 vs 500 vs 500 mM sucrose<br>1% agarose mixture, 0 vs 200 vs 200 mM sucrose<br>1% agarose mixture vs 2% agarose mixture vs 2% agarose mixture |
| 2 | a-h | wheel | 1% agarose mixture, 0 vs 0 vs 0 mM sucrose |
| 3 | a | free behavior choice | 1% agarose mixture, 0 vs 500 mM sucrose |
| 3 | b | free behavior choice | 1% agarose mixture, 0 vs 500 mM sucrose<br>1% agarose mixture, 0 vs 200 mM sucrose<br>1% agarose mixture, 200 vs 500 mM sucrose |
| 3 | c | free behavior choice | 1% agarose mixture, 0 vs 500 mM sucrose<br>1% agarose mixture, 0 vs 200 mM sucrose<br>1% agarose mixture, 200 vs 500 mM sucrose<br>1% agarose mixture, 0 vs 0 mM sucrose<br>1% agarose mixture, 200 vs 200 mM sucrose<br>1% agarose mixture, 500 vs 500 mM sucrose |
| 3 | d-f | free behavior choice | 1% agarose mixture, 0 vs 500 mM sucrose |
| 3 | g | free behavior choice | 1% agarose mixture, 0 vs 200 mM sucrose |
| 3 | h | free behavior choice | 1% agarose mixture, 200 vs 500 mM sucrose |
| 4 | b-c | wheel | 1% agarose mixture, 0 vs 0 vs 500 mM sucrose<br>1% agarose mixture, 0 vs 500 vs 500 mM sucrose |
| 4 | de | wheel | 1% agarose mixture, 0 vs 0 vs 500 mM sucrose<br>1% agarose mixture, 0 vs 500 vs 500 mM sucrose<br>1% agarose mixture, 0 vs 200 vs 500 mM sucrose |
| 5 | a-c | free behavior choice | 1% agarose mixture, 0 vs 200 mM sucrose |
| 5 | d | wheel | 1% agarose mixture, 0 vs 500 vs 500 mM sucrose |
| 5 | e-i | free behavior choice | 1% agarose mixture, 0 vs 200 mM sucrose |
| Extended Data Fig 2 | a-e | free behavior sloped ceiling | 1% agarose mixture, 0 mM sucrose |
| Extended Data Fig 3 | d | free behavior choice | 1% agarose mixture, 0 vs 200 mM sucrose |
| Extended Data Fig 3 | d (last bar) | free behavior choice<br>with bottom 2 layers printed<br>in VisiJet M3 Crystal material | 1% agarose mixture, 0 vs 200 mM sucrose |
| Extended Data Fig 3 | e | free behavior choice | 1% agarose mixture, 0 vs 200 mM sucrose |
| Extended Data Fig 4 | a | wheel | 1% agarose mixture, 0 vs 500 vs 200 mM sucrose |
| Extended Data Fig 4 | b-l | wheel | 1% agarose mixture, 0 vs 0 vs 500 mM sucrose<br>1% agarose mixture, 0 vs 500 vs 500 mM sucrose<br>1% agarose mixture, 0 vs 200 vs 500 mM sucrose<br>1% agarose mixture, 0 vs 200 vs 200 mM sucrose<br>1% agarose mixture vs 2% agarose mixture vs 2% agarose mixture |
| Extended Data Fig 6 | a | wheel | 1% agarose mixture, 0 vs 0 vs 0 mM sucrose |
| Extended Data Fig 6 | b | wheel | 1% agarose mixture, 0 vs 0 vs 500 mM sucrose<br>1% agarose mixture, 0 vs 500 vs 500 mM sucrose<br>1% agarose mixture, 0 vs 200 vs 500 mM sucrose<br>1% agarose mixture, 0 vs 200 vs 200 mM sucrose<br>1% agarose mixture vs 2% agarose mixture vs 2% agarose mixture |
| Extended Data Fig 6 | c | wheel | 1% agarose mixture, 0 vs 0 vs 500 mM sucrose<br>1% agarose mixture, 0 vs 500 vs 500 mM sucrose<br>1% agarose mixture, 0 vs 200 vs 500 mM sucrose<br>1% agarose mixture, 0 vs 500 vs 200 mM sucrose<br>1% agarose mixture, 0 vs 200 vs 200 mM sucrose<br>1% agarose mixture, 0 vs 0 vs 0 mM sucrose<br>1% agarose mixture, 200 vs 200 vs 500 mM sucrose<br>1% agarose mixture vs 2% agarose mixture vs 2% agarose mixture |
| Extended Data Fig 6 | d | wheel | 1% agarose mixture, 0 vs 0 vs 500 mM sucrose<br>1% agarose mixture, 0 vs 500 vs 500 mM sucrose<br>1% agarose mixture, 0 vs 200 vs 500 mM sucrose<br>1% agarose mixture, 0 vs 500 vs 200 mM sucrose<br>1% agarose mixture, 0 vs 0 vs 0 mM sucrose<br>1% agarose mixture, 200 vs 200 vs 500 mM sucrose<br>1% agarose mixture vs 2% agarose mixture vs 2% agarose mixture |
| Extended Data Fig 7 | a | wheel | 1% agarose mixture, 0 vs 0 vs 500 mM sucrose<br>1% agarose mixture, 0 vs 500 vs 500 mM sucrose<br>1% agarose mixture, 0 vs 200 vs 500 mM sucrose<br>1% agarose mixture, 0 vs 500 vs 200 mM sucrose<br>1% agarose mixture, 0 vs 200 vs 200 mM sucrose<br>1% agarose mixture, 0 vs 0 vs 0 mM sucrose<br>1% agarose mixture, 200 vs 200 vs 500 mM sucrose<br>1% agarose mixture vs 2% agarose mixture vs 2% agarose mixture |
| Extended Data Fig 7 | b | wheel | 1% agarose mixture, 0 vs 0 vs 500 mM sucrose |
| Extended Data Fig 7 | cd (1st 4 cross<br>-correlations) | wheel | 1% agarose mixture, 0 vs 0 vs 500 mM sucrose<br>1% agarose mixture, 0 vs 500 vs 500 mM sucrose<br>1% agarose mixture, 0 vs 200 vs 500 mM sucrose<br>1% agarose mixture, 0 vs 500 vs 200 mM sucrose<br>1% agarose mixture, 0 vs 200 vs 200 mM sucrose<br>1% agarose mixture, 0 vs 0 vs 0 mM sucrose<br>1% agarose mixture, 200 vs 200 vs 500 mM sucrose<br>1% agarose mixture vs 2% agarose mixture vs 2% agarose mixture |
| Extended Data Fig 7 | cd (5th cross<br>-correlation) | wheel | 1% agarose mixture, 0 vs 0 vs 500 mM sucrose<br>1% agarose mixture, 0 vs 500 vs 500 mM sucrose |
| Extended Data Fig 7 | e | wheel | 1% agarose mixture, 0 vs 0 vs 500 mM sucrose<br>1% agarose mixture, 0 vs 500 vs 500 mM sucrose<br>1% agarose mixture, 0 vs 200 vs 500 mM sucrose<br>1% agarose mixture, 0 vs 500 vs 200 mM sucrose<br>1% agarose mixture, 0 vs 200 vs 200 mM sucrose<br>1% agarose mixture, 0 vs 0 vs 0 mM sucrose<br>1% agarose mixture, 200 vs 200 vs 500 mM sucrose<br>1% agarose mixture vs 2% agarose mixture vs 2% agarose mixture |
| Extended Data Fig 8 | a-i | wheel | 1% agarose mixture, 0 vs 0 vs 0 mM sucrose |
| Extended Data Fig 9 | b | free behavior choice | 1% agarose mixture, 0 vs 500 mM sucrose |
| Extended Data Fig 9 | c | free behavior choice | 1% agarose mixture, 0 vs 200 mM sucrose |
| Extended Data Fig 9 | d | free behavior choice | 1% agarose mixture, 200 vs 500 mM sucrose |
| Extended Data Fig 10 | a-c | free behavior choice | 1% agarose mixture, 0 vs 0 mM sucrose<br>1% agarose mixture, 200 vs 200 mM sucrose<br>1% agarose mixture, 500 vs 500 mM sucrose |
| Extended Data Fig 11 | a-c | wheel | 1% agarose mixture, 0 vs 0 vs 500 mM sucrose<br>1% agarose mixture, 0 vs 500 vs 500 mM sucrose |
| Extended Data Fig 12 | a | wheel | 1% agarose mixture, 0 vs 0 vs 500 mM sucrose<br>1% agarose mixture, 0 vs 500 vs 500 mM sucrose |
| Extended Data Fig 12 | b | wheel | 1% agarose mixture, 0 vs 0 vs 500 mM sucrose |
| Extended Data Fig 12 | c-e | wheel | 1% agarose mixture, 0 vs 0 vs 500 mM sucrose<br>1% agarose mixture, 0 vs 500 vs 500 mM sucrose |
| Extended Data Fig 13 | a,d | wheel | 1% agarose mixture, 0 vs 0 vs 500 mM sucrose<br>1% agarose mixture, 0 vs 500 vs 500 mM sucrose |
| Extended Data Fig 13 | b,e | wheel | 1% agarose mixture, 200 vs 200 vs 500 mM sucrose |
| Extended Data Fig 13 | c,f | wheel | 1% agarose mixture, 0 vs 200 vs 200 mM sucrose |
| Extended Data Fig 14 | a-c | free behavior choice | 1% agarose mixture, 0 vs 200 mM sucrose |
| Extended Data Fig 15 | a | wheel | 1% agarose mixture, 0 vs 500 vs 500 mM sucrose |
| Extended Data Fig 15 | b-c | free behavior choice | 1% agarose mixture, 0 vs 200 mM sucrose |
| Supplementary Video 1 |  | free behavior sloped ceiling | 1% agarose mixture, 0 mM sucrose |
| Supplementary Video 2 |  | free behavior sloped ceiling | 1% agarose mixture, 0 mM sucrose |
| Supplementary Video 3 |  | wheel | 1% agarose mixture, 0 vs 0 vs 500 mM sucrose |
| Supplementary Video 4 |  | wheel | 1% agarose mixture, 0 vs 0 vs 500 mM sucrose |
| Supplementary Video 5 |  | wheel | 1% agarose mixture, 0 vs 0 vs 0 mM sucrose |
| Supplementary Video 6 |  | free behavior choice | 1% agarose mixture, 0 vs 500 mM sucrose |

### Supplemental Videos

**Supplementary Video 1.** Individual egg-laying event of a fly expressing GCaMP3 in eggs. An egg-laying event with the behavioral sequence occurring in a localized space was chosen so that it could be magnified. Video is compressed and played back at 5X speed.

**Supplementary Video 2.** Three consecutive egg-laying events of a wild type fly in a sloped ceiling egg-laying chamber (as in Extended Data Fig. 2a). Traces below movie are smoothed with a 5 s boxcar filter. Pink line is overlaid on fly to indicate the neck to ovipositor length and is only drawn in frames that passed the criteria described in the Methods. Video is compressed and played back at 1X speed.

**Supplementary Video 3.** Egg-laying event during two-photon imaging in a tethered fly expressing GCaMP7f in oviDNs (as in Fig. 1g left). ΔF/F and brain images are smoothed with a 2 s boxcar filter. Video is compressed and played back at 5X speed.

**Supplementary Video 4.** Egg-laying event during two-photon imaging in a tethered fly expressing GCaMP7f in oviDNs (as in Fig. 1g right). ΔF/F and brain images are smoothed with a 2 s boxcar filter. Video is compressed and played back at 5X speed.

**Supplementary Video 5.** Optogenetically stimulated egg-laying event and abdomen bends in a tethered fly expressing GCaMP7f and CsChrimson in oviDNs (as in Fig. 2a left). CsChrimson stimulations here were manually triggered. An orange dot in close-up of fly and orange line in ‘fly position’ trace both indicate stimulation periods. ΔF/F and brain images are smoothed with a 2 s boxcar filter. Video is compressed and played back at 5X speed.

**Supplementary Video 6.** Wild type flies in a high-throughput 0 (bottom) vs. 500 (top) mM egg-laying chamber. Video is compressed and sped up 60X by displaying only every 60^th^ frame. The analyzed movies for figures included all the frames and were therefore much smoother.

## Methods

### Flies

Flies were reared on a standard corn-meal medium at 25°C, ambient humidity, and a 12 h/12 h light/dark cycle unless otherwise noted. Genotypes and conditions for each experiment are described in Supplementary Table 1 and Supplementary Table 2, respectively. Supplementary Table 1 also lists the source of each genotype.

### Egg-laying chamber with sloped ceiling

We designed a new chamber for imaging egg laying in freely walking flies, which enforced flies to remain in a tarsi-down body posture over the agarose at all times. The flies could not tilt their bodies in this chamber and thus they could not walk on the side walls or ceiling. This constraint meant that the flies’ bodies were always in the same general orientation, parallel to the imaging plane, throughout, allowing for quantitative measurements of postural parameters.

Chambers were made by sandwiching and tightly screwing layers of acrylic and 3D-printed plastic and then placing a glass ceiling (Extended Data Fig. 1a). The acrylic layers were laser-cut (VLS6.60, Universal Laser Systems). The side-wall layer was 3D-printed using the VisiJet M3 Crystal plastic material (Projet 3510 HD Plus, 3D Systems). The glass was treated with Sigmacote (Sigma-Aldrich) to make it slippery to a fly’s tarsi––preventing any walking on the ceiling^65^. Glass was re-treated with Sigmacote after ∼10 uses. The 3D-printed spacer layer incorporated a sloped edge that kept the fly completely parallel to the imaging plane by preventing access to the side of the chamber (Extended Data Fig. 1a). This allowed for quantitative measurements of body posture––e.g., the distance from neck to ovipositor––which would be distorted if the fly were able to tilt. The sloped ceiling design was inspired by a sloped floor plastic chamber^65^. A sloped floor does allow the fly to tilt and thus was not suitable for our application.

Chambers were used multiple times and washed prior to each use. Chambers were assembled with only the two bottom layers and then cooled at 4°C. Fresh substrate containing 1% agarose (SeaKem LE Agarose, Lonza), 0.8% acetic acid, and 1.6% ethanol was pipetted to completely fill the well ∼5 hrs prior to each assay. Careful pipetting with only the two bottom layers assembled was critical to forming a flat layer of agarose––preventing the formation of a meniscus, which would allow the fly to tilt. Acetic acid and ethanol were included to help simulate a rotten fruit and generally promote egg laying^9^. After the agarose solution was solidified (∼1 hr), the chamber was fully assembled, minus the glass ceiling, and equilibrated at room temperature.

Females were separated on their day of eclosion and group housed in vials. At age 3 to 6 days, ∼20 females were exposed to ∼20 Canton-S males in an empty bottle with wet yeast paste and a Kimwipe (Kimberly-Clark) soaked with 2 ml of water. The wet yeast paste was applied to the side of the bottle and was composed of 1-gram dry yeast (Fleischmann’s) and 1.5 ml of 4.25 mM putrescine dihydrochloride in water. This treatment allowed females to mate and caused them to accumulate many eggs. Flies fed with yeast^9, 66^ or putricine^20^ increase the number of eggs they develop. These eggs are retained by the flies during the treatment period because they lack a soft medium for egg deposition^9^. After ∼24 hrs, individual gravid females were placed into chambers under gentle cold anesthesia from which they typically recovered within 30 s. The fly with the least ability to tilt (of 6 flies in individual chambers) was chosen to be imaged for a few hours in near-complete darkness (under a shroud) at ∼24°C and 40-60% humidity.

For imaging of eggs inside the body, a 470 nm LED (pE-100, CoolLED) filtered twice (OD4 475 nm and OD4 500 nm shortpass, Edumund Optics) provided excitation light at 30 µW/mm. This excitation light arrived at the fly from below, after first passing through the agarose substrate. Videos were recorded at 10 fps with a 100 ms exposure time per frame, using an ORCA-Fusion C14440-20UP camera (Hamamatsu) equipped with a 15.5-20.4 mm Varifocal Lens (Computar) and two 510 nm longpass filters (Chroma). Representative stills (Fig. 1b) or videos (Extended Data Video 1) were selected.

For imaging body posture, 850 nm LEDs illuminated the arena from above, through a white acrylic diffuser (1 µW/mm^2^ at the fly). Videos were recorded at 25 fps using a GS3-U3-41C6NIR-C Grasshopper camera (FLIR) equipped with a 15.5-20.4 mm Varifocal Lens and a 780 nm longpass filter (MidOpt). DeepLabCut^67^ was used for offline tracking of body parts including the neck and ovipositor. DeepLabCut models were iteratively fine-tuned by identifying poorly tracked frames in iteration *i* and adding them to the training dataset for iteration *i+1*. A total of 1568 training frames were manually annotated. DeepLabCut output coordinates were filtered by setting coordinates to NaN if (1) the probability score was less than 0.95; or (2) the body part jumped more than an empirically determined distance in consecutive frames. ‘Ovulation start’ was defined as the first frame in which the abdomen appeared to begin the elongation process. ‘Search start’ was manually annotated as the first frame in which the abdomen returned to a stable neutral posture after ovulation. ‘Abdomen bend complete’ was manually annotated as the frame in which the bend to lay an egg was completed (abdomen maximally deflected). Identifying the frame in which the abdomen bend was completed was much easier than attempting to identify when then abdomen bend was initiated. Note that although flies bend their abdomen to deposit an egg, flies also bend their abdomen during our experiments for other reasons. ‘Egg deposited’ was manually annotated (with some computer assistance) as the first frame in which half the egg was visible (emerging from the ovipositor).

### High-throughput egg-laying choice chamber

We designed a new chamber for studying egg-laying choice behavior with high-throughput. This chamber ensured the fly was nearly always in contact with an agarose egg-laying substrate option. The substrate the fly was on could be unambiguously defined by the fly’s y-position and orientation. In previous egg-laying choice studies^10, 19, 68^, flies could walk on the side walls or ceiling and yet were assigned to a substrate beneath them during tracking, which makes it very hard to determine how previous substrate experiences influence the decision to lay an egg.

Chambers were made by sandwiching and tightly screwing layers of acrylic or Delrin plastic and then placing a glass ceiling (Extended Data Fig. 1b). Acrylic and Delrin plastic were laser-cut. The glass was treated with Sigmacote.

Chambers were used multiple times and washed prior to each use. Chambers were assembled without the glass ceiling and cooled at 4°C. 1 ml of fresh substrate containing 1% agarose, 0.8% acetic acid, and 1.6% ethanol was pipetted to fill the acrylic well and form a meniscus with the Delrin plastic spacer ∼5 hrs prior to each assay. The meniscus ensured that the fly could not walk directly on the side (Delrin plastic) of the chamber and was inspired by plastic chambers with a sloped floor^65^. Quantitative measurements of body posture were not possible because flies could tilt by walking on the meniscus. Sucrose containing substrates were supplemented with the appropriate amount of sucrose. Acetic acid and ethanol were uniformly distributed in all substrates. After the agarose solution was solidified (∼1 hr), the chamber was equilibrated at room temperature.

These egg-laying chambers and assay protocols were specifically designed to minimize the following confounds: (1) diffusion between substrate islands; (2) visual landmarks; (3) fly-to-fly communication; (4) olfactory landmarks; (5) temperature and humidity fluctuations; and (6) variability in fly rearing. Diffusion was minimized by a ∼2.5 mm wide barrier between the substrate islands and by loading the agarose at 4°C. Visual cues were minimized by conducting the assay in near-complete darkness. 850 nm illumination, to which the fly visual system has no measurable sensitivity^69–71^, was provided from below for tracking (1 µW/mm^2^ at the agarose beneath the fly). Fly-to-fly communication was minimized by assaying individual flies in isolated chambers separated by an opaque Delrin plastic spacer. Olfactory landmarks were minimized by using a non-volatile compound, sucrose, as the sole varying variable. Temperature and humidity were kept constant by conducting experiments in an environmental room (24°C with 40-60% humidity). Air exchange was made possible by four small ventilation holes in each barrier. Variability in fly rearing was minimized by controlling age, mating status, food history, and circadian time.

Females and males were separated on their day of eclosion and group housed in vials. At age 3 to 6 days at zeitgeber time (ZT) 6 (i.e., 6 hours after lights on), ∼20 females were exposed to ∼20 Canton-S males in an empty bottle with only wet yeast paste and a Kimwipe soaked with 2 ml of water. Putrescine was not added to the yeast paste in these experiments because we could always test sufficient flies wherein the additional eggs provided by feeding putrescine was not required. On the next day at ZT 8, individual females were placed into egg-laying chambers under gentle cold anesthesia. Images were acquired at 2 fps with a FMVU-03MTM-CS Firefly or FL3-U3-13Y3M-C Flea3 camera (FLIR) equipped with either a LM12HC (Kowa), HF12.5SA-1 (Fujinon), or CF12.5HA-1 (Fujinon) lens and a 780 nm longpass filter. The x-y position and orientation of each fly was extracted offline using Ctrax^72^. We assigned a fly’s current substrate depending on whether its centroid was above or below the midline of the acrylic barrier. This simplification was appropriate because the ∼2.5 mm acrylic barrier (a fly is ∼2.5 mm long) practically prevented a fly from standing on both substrates simultaneously and a Canton-S fly spent only 1.5% of its time in an orientation where all tarsi were likely to be on the barrier. Note that flies do not lay eggs on the plastic barrier (or any plastic used in this study) because it is too hard. ‘Egg deposition’ was manually annotated (with some computer assistance) as the first frame in which half the egg was visible (emerging from the ovipositor). Annotations by an individual human annotator or across multiple human annotators were reproducible to ± 4 frames or ± 2 s. The ‘search period’ was determined as described in the section ‘Automated estimation of search…’.

For Kir2.1* or Kir2.1*Mut experiments, we expressed a GAL80^ts^ transgene in all cells (with the tubulin promoter)^73^ during development so as to minimize transcription of the Kir transgenes days before assaying egg-laying behavior. At 18°C, GAL80^ts^ masks the transcription activation domain of GAL4 preventing transcription of the GAL4-UAS controlled transgene. We could remove the GAL80 block on Kir expression by increasing the flies’ temperature for ∼1 day prior to our egg-laying assays. Specifically, for these experiments: (1) flies were reared at 18°C; (2) at ZT 6 flies were shifted to 31°C for induction of Kir2.1* or Kir2.1*Mut transgene expression; and (3) the next day at ZT 5 (23 hours later), flies were returned to 18°C. Egg-laying assays were performed at ZT 8 at 24°C.

For GtACR1^35, 74^ experiments, 567 nm light was provided from above (29 µW/mm^2^ at the fly) (Rebel Tri Star LEDs, LuxeonStarLEDs). Controls for genotype were siblings of experimental flies that were treated identically except no light was provided from above. Controls for light were flies ‘expressing’ GtACR1 with an empty split or empty GAL4 driver. Additional controls for light with twice the intensity (57 µW/mm^2^) provided ample assurance that light alone was not preventing egg laying (data not shown). 567 nm light was chosen to minimize overlap with the sensitivity of the fly visual system^69–71^ while still stimulating GtACR1.

### Construction of Kir2.1* and Kir2.1*Mut flies

We screened a variety of reagents to find a way to gently hyperpolarize oviDNs. We serendipitously identified a Kir2.1* that, as inferred from our behavioral experiments (Fig. 5c), hyperpolarizes oviDNs more gently than the Kir2.1 traditionally used in flies^34, 75, 76^ (Fig. 5a). A matched control, Kir2.1*Mut, that has a non-conducting channel facilitates quantitative comparisons. In theory, a ‘low’ level of light stimulating GtACR1/2 should also be able to gently hyperpolarize neurons^35, 74^. However, (1) identical illumination of each fly was hard for us to practically achieve because multiple chambers were spread across a large area that needed to be evenly illuminated and flies can rotate their heads exposing neurons to more or less stimulation as a result; (2) illumination is hard to match between free behavior and tethered electrophysiology preparations; (3) a non-conducting control for GtACR1/2 does not currently exist, so separate controls for genotype and light are necessary; and (4) egg-laying assays would no longer be in the dark.

Kir2.1* and Kir2.1*Mut sequences were taken from a previous study in mice^36^. Briefly, Kir2.1* and Kir2.1*Mut are mouse wild type Kir2.1s (KCNJ2) – with two mutations (Kir2.1*: E224G, Y242F) or five mutations (Kir2.1*Mut: E224G, Y242F, G144A, Y145A, G146A). Both transgenes were fused at their C-terminals with a T2A sequence to a tdTomato. To port these constructs into *Drosophila,* they were inserted between the Xba1 and Not1 sites of pJFRC81^77^ and introduced into the *attP40* landing site by ΦC31 integrase-mediated transgenesis (transgenic fly lines were generated by BestGene). Kir2.1* and Kir2.1*Mut transgenes differ in protein sequence––and possibly in other ways (e.g., transcription and translation)–––from the wild type human Kir2.1 (KCNJ2) transgenes traditionally used to hyperpolarize neurons in flies^34, 75, 76^. Previous in vivo fly electrophysiology of central brain and visual system neurons expressing traditional human Kir2.1^78, 79^ transgenes showed larger hyperpolarization than the ∼14 mV hyperpolarizations observed here with Kir2.1* (Fig. 5d).

### Automated estimation of search period in free behavior high-throughput choice chambers

Since we did not have a high-resolution view of the abdomen in our high-throughput choice chambers (Extended Data Fig. 1c), we used locomotor speed as a proxy for search onset (Extended Data Fig. 2bc) and egg deposition as a proxy for abdomen bending to lay an egg (Fig. 1c). The end of the search period was the annotated moment of egg deposition (rather than the abdomen bend to lay the egg). The start of the search period was determined for each egg by smoothing the locomotor speed trace, prior to egg deposition, with an 18.5 s boxcar filter and identifying the first frame where this smoothed signal dropped below 0.1 mm/s. The minimum search duration was 18.5 s due to the length of the boxcar filter. These parameters were determined empirically to yield search onset times consistent with visual inspection of the data.

### Calculation of egg-laying rates as a function of time since the last substrate transition in free behavior choice chambers

Egg-laying rates as a function of time (Figure 3f-h) were calculated as follows. Data from all flies tested in a given chamber type were combined prior to any calculations. First, we iterated through each time bin denoted on the x-axis and, for each bin, we counted the number of egg-deposition events that were assigned to that bin, #_eggs_(bin). Second, we iterated through the same time bins and counted the number of video frames in which flies were assigned to that bin, #_frames_(bin), during an egg-laying search period. Third, we iterated through the same time bins and counted the number of times flies changed assignment into that bin, #_visits_(bin), during an egg-laying search period (i.e., we did not keep incrementing the “visits” counter if the fly remained in a time bin from one frame to the next).

To get the mean egg-laying rate, we calculated #_eggs_/#_frames_ for each bin. Since videos were recorded at 2 fps, we multiplied the value for each bin by 120 to convert to units of eggs/minute.

To get the confidence interval per bin, we used the Clopper-Pearson method (‘exact’ binomial confidence interval) to calculate the 90% confidence interval for #_eggs_/#_visits_ for each bin. We then converted the confidence interval for each bin to units of eggs/minute by multiplying by 120*#_visits_/#_frames_ per bin. Note the confidence interval cannot be directly calculated from #_eggs_/#_frames_ because then the confidence interval would be dependent on the video framerate.

For these rate functions, search periods with duration less than 30 s were set to 30 s. This prevented very short search periods from introducing fluctuations in the rate curves (by contributing to the numerator and not contributing much to the denominator). As such, rate curves varied less from replicate-to-replicate or condition-to-condition. Note that search periods already had a minimum duration of 18.5 s as automatically generated by the search period calculation (Methods). Analyzing the data with no minimum search duration (or with other definitions of the search period) did not change any of the stated conclusions (data shown in a second manuscript *in preparation*). Binning the x-axis differently also did not qualitatively change any of our stated conclusions. Rate functions start off with low rates after a transition at least partially because flies do not lay eggs on the plastic barrier between substrates (Extended Data Figure 10a-b) and because flies are, by definition, walking (and not pausing to deposit an egg) during a transition (Extended Data Figure 10c).

### Design of egg-laying wheel and setup under microscope

We designed a wheel upon which tethered flies walked and laid eggs on agarose-based egg-laying substrates. The design was optimized to maximize a fly’s ability to lay eggs and rotate the wheel.

The wheels were 3D-printed from VisiJet M3 Crystal plastic using a Projet 3510 HD Plus 3D printer (Extended Data Fig. 3a). A pivot (N-1D, Swiss Jewel) was press fit through the center hole and never removed. Wheels were washed prior to each use. Three wells were available for loading the same or different agarose-based substrates. Each well was separated by a 1 mm barrier. Wheels were loaded with fresh agarose substrates (as prepared for free behavior choice chambers) using a 3D-printed agarose-injecting mold (VisiJet M3 Crystal material) that was cooled on ice (Extended Data Fig. 3b). Food coloring (HY-TOP assorted food coloring) was added at a dilution of 1:10,000 to the agarose solution prior to loading so wheel quality could be visualized. Wheels with any mixing between wells were discarded. Food coloring at 2.5 times this concentration or the presence of VisiJet M3 Crystal material did not affect choice in free behavior control experiments (Extended Data Fig. 3d). After the agarose was solidified, the wheel and pivot were suspended between two spring loaded bearings (VS-30, Swiss Jewel) that were threaded into clear acrylic that was press fit into a 3D-printed base (UMA-90 material printed on a Carbon DLS, Protolabs) (Extended Data Fig. 3c). This wheel assembly was stored in a custom humidification chamber to prevent the thin layer of agarose from drying and to allow the wheels to equilibrate to room temperature. Wheels were used within 2 hrs of preparation. When ready, a wheel assembly was secured in a small custom humidification chamber (∼90% humidity) that sits under the microscope objective attached to a micromanipulator. The wheel-pivot combinations used in this study had weight 87.9 ± 0.3 mg (mean ± std) without agarose and 146.4 ± 0.8 mg with agarose. For reference, a single gravid female weights ∼1.4 mg and a typical foam ball used for fly walking experiments^80, 81^ weighs 40 to 46 mg. Most of the wheel’s weight is due to the agarose and the wells needed to hold it. A variety of lighter and less-prone-to-evaporation synthetic materials were screened in free behavior assays, but egg laying was suppressed on all of them.

The fly was viewed by two CM3-U3-13Y3M Chameleon cameras (from the sides) and one FMVU-03MTM-CS Firefly camera (from the front) (FLIR). Two 850 nm LEDs from the front-left and front-right illuminated the fly at 5 µW/mm^2^. Cameras were equipped with a 15.5-20.4 mm Varifocal Lens and either a 900 nm shortpass (Thorlabs) or 875 nm shortpass (Edmund Optics) filter to dampen visibility of the 925 nm two-photon excitation light. Cameras had an exposure time of 16 ms and were triggered synchronously using a single external trigger source at 25 fps (Arduino Uno, Arduino). A side facing camera recorded the fly and a single dot painted on the wheel. The dot was painted in a consistent location on the wheel that was defined by an embossed 3D-printed feature. The dot was tracked using DeepLabCut (1109 training frames, with training and filtering like the free behavior DeepLabCut model).

The dot position was converted to wheel degrees by fitting the set of all dot positions to a circle and then computing a wheel angle for each frame. A single frame in which the fly’s centroid straddled the dot was used to convert the wheel angle to the fly’s position on the wheel. This alignment consistently meant that the fly’s neck was situated on the plastic-to-next-substrate boundary during a detected substrate transition. A second side facing camera was used for a close-up view of the fly’s body. DeepLabCut was used to track body parts including the neck, ovipositor, and tip of the proboscis (2259 training frames, with training and filtering like the free behavior DeepLabCut model). ‘Normalized length’ was calculated by subtracting the x-coordinates of the neck and ovipositor in each frame and dividing by the median of this value for each recording (Fig. 1i). An absolute reference for length was not easy to include in the camera frame and therefore distance was measured about the median. For reference, the median length in free behavior was ∼2.35 mm (Extended Data Fig. 2b-e). This metric for length was used since it can quantify both an elongated and a bent abdomen and is similar to the neck to ovipositor length measured in free behavior. Despite the similarity with free behavior length, we noticed, on average, a slight difference in the signature of abdomen bends (Extended Data Fig. 2d compared to Fig. 1l) possibly due to the curvature of the wheel. The ‘body angle (°)’ was the angle between the neck and ovipositor (Fig. 2i). Larger angles indicated a more bent abdomen. Although a fly must bend its abdomen to lay an egg, the magnitude of a physiologically relevant deflection of body angle (as measured in degrees) is not that large (Fig. 2i). ‘Normalized neck to proboscis length’ was calculated by determining the Euclidean distance between the tip of the proboscis and the neck in each frame and dividing by the median of this value for each recording. This underestimated the true deflection of the proboscis because the proboscis does not start at the neck. The neck was used as an origin point because it was easy to robustly track. The position of each tarsus was not able to be tracked, and the tarsi on the opposite side of the camera were often occluded by the body. A front facing camera was used to align the fly on the center of the wheel width. The body posture slightly varied from fly-to-fly due to slight differences in tethering. To achieve egg laying, it was very important to position the fly at a point on the wheel circumference, and at a vertical distance from wheel, that maximized perpendicular contact of the ovipositor to the substrate when the abdomen was bent while still allowing the fly to walk on the wheel. In some cases, flies had to be positioned close to the wheel which, unfortunately, decreased the dynamic range of abdomen bending. 104 flies were imaged, to collect the data displayed in Figure 1h. The majority of flies did not lay eggs because, among other considerations, it often takes flies several hours to start laying a “clutch” of eggs (even when freely behaving in a chamber) and we could not image conveniently for 18 hours to wait for a clutch to start.

Moments of distinct behaviors (as in Fig. 2h and Extended Data Fig. 4b) were annotated manually by inspecting the behavior videos while being blind to any neural signals (ΔF/F). ‘Ovulation start’ was defined as the first frame in which the abdomen appeared to begin the elongation process. ‘Abdomen at its longest’ was the frame in which the abdomen was maximally stretched. ‘Abdomen scrunch start’ was the first frame in which the abdomen assumed a stable scrunched position. ‘Search start’ was defined as the first frame in which the abdomen returned to a stable neutral posture after ovulation. ‘Abdomen bend complete’ was defined as the frame in which the first bend prior to egg laying was complete (abdomen maximally deflected). ‘Egg deposited’ was defined as the frame in which half the egg was visible. ‘Ovipositor cleaned’ was defined as the frame in which the first abdomen bend following egg laying was complete.

For CsChrimson^32^ optogenetics experiments, a 660 nm LED coupled to a 1 mm wide lightguide (M660F1 and M35L01, Thorlabs) was focused on the front, midpoint of the fly’s head using a lens set (MAP10100100-A, Thorlabs). A wavelength at the very tail end of the sensitivity of the fly visual system^69–71^ that could still stimulate CsChrimson was used to minimize light-related confounds. Two shortpass filters––OD4 550 nm and OD 4 575 nm (Edmund Optics)––minimized the ability of the LED light to enter the two-photon detector path, which collected the GCaMP signal. The incident area of the LED was adjusted to be wide enough (∼3 mm diameter) to cover the whole front of the fly such that all CsChrimson expressing oviDN cell bodies and neurites in the brain (and likely in the thorax) could be stimulated. The LED intensity was controlled by adjusting the duty cycle of a 490 Hz PWM signal (Arduino Uno, Arduino) that was fed into an LED driver (T-Cube, Thorlabs). The CsChrimson stimulation intensity for Figure 2a-e was 641 µW/mm^2^. For Figure 2f-h intensities were 641, 159, 148 and 136 µW/mm^2^.

### Treatment of flies for tethered experiments

Females and males were collected on their day of eclosion and group housed together in standard corn-meal medium vials supplemented with 2.5 mM putrescine dihydrochloride and wet yeast paste. The wet yeast paste was applied to the side of the vial and was composed of 1-gram dry yeast and 1.5 ml of 4.25 mM putrescine dihydrochloride in water. At age ∼5-6 days females were gravid because larvae occupied the corn-meal medium and there was no additional room to deposit eggs. This treatment was more convenient than the treatment used in free behavior choice experiments and was inspired by separate aspects of two studies^10, 20^. Free behavior controls indicated that this treatment increased the number of eggs a fly laid without affecting choice behavior (Extended Data Fig. 3e).

For CsChrimson optogenetics experiments, flies were treated as above, but were kept in low white light (∼3 nW/mm^2^ measured at 660 nm) from egg to adulthood. At age ∼5-6 days, ∼20 females were exposed to ∼20 Canton-S males in an empty bottle with only wet yeast paste and a Kimwipe soaked with 2 ml of 200 µM all-trans-retinal in water. The wet yeast paste was applied to the side of the bottle and was composed of 1-gram dry yeast with 1.5 ml of 4.25 mM putrescine dihydrochloride and 200 µM all-trans-retinal in water. Flies were tethered ∼24 hours later. Flies for CsChrimson control experiments were always treated identically to CsChrimson expressing flies.

Flies were anesthetized at ∼4°C and tethered to a custom holder^82^, except where the back wall of the pyramid leading up to the fly was tilted at an angle instead of rising at 90° so as to allow more light from the brain to reach the objective^81^ (Fig. 1d). The head was pitched forward during tethering to provide a view of the oviDN cell bodies. For electrophysiology, the head was inserted deeper into the holder for unobstructed access to the oviDNs with electrodes. Flies were attached to the holder with blue-light-cured glue (Bondic). The proboscis was gently extended, and the dorsal rostrum was glued to the head capsule. This prevented brain movement associated with proboscis extension while still allowing proboscis extension to be measured (albeit with a smaller dynamic range than natural proboscis extension). Extracellular saline solution was added to the holder well (bath) and a window was cut in the cuticle to expose the posterior side of the brain. The holder was stabilized with magnets above the egg-laying wheel inside a small custom humidification chamber.

Our extracellular saline^83^ was composed of 103 mM NaCl, 3 mM KCl, 5 mM N-Tris(hydroxymethyl) methyl-2-aminoethanesulfonic acid, 10 mM trehalose, 10 mM glucose, 2 mM sucrose, 26 mM NaHCO_3_, 1 mM NaH_2_PO_4_, 1.5 mM CaCl_2_, and 4 mM MgCl_2_. The osmolarity was 280 ± 5 mOsm, and the pH was 7.3-7.4 when bubbled with 95% O_2_ / 5% CO_2_. The temperature of the bath was set to ∼19-21°C by flowing fresh saline through a Peltier device with feedback from a thermistor in the bath (Warner Instruments).

### Calcium imaging

We used a two-photon microscope with a movable objective (Ultima IV, Bruker) and custom stage (Thorlabs, Siskiyou). The microscope was controlled by Prairie View software (Bruker) and was enclosed by a black shroud. A Chameleon Ultra II Ti:Sapphire femtosecond pulse laser (Coherent) filtered by a 715 nm longpass filter (Semrock) provided 925 nm two-photon excitation. Emission light from the brain was collected by a 16x/0.80 NA objective (16X W CFI75 LWD, Nikon), split by a 565 nm dichroic, and filtered by a 490 to 560 nm bandpass filter (Chroma) prior to entering GaAsP detectors (Hamamatsu). For CsChrimson optogenetics experiments, the emission light was split by a 525 nm dichroic and filtered by both a 490 to 510 nm bandpass and a 480 to 520 nm bandpass filter (Chroma) to prevent optogenetic stimulation light from entering the detector. A Piezo motor was used for volumetric scanning.

A range of optical zooms, z-slice number, z-slice separation, fields of view, laser powers (6 to 30 mW at the specimen), and frame rates (mean of 1.84 Hz) were used over the course of experiments on oviDN dynamics. Individual data traces were inspected by eye and the reported results were robust to the range of parameters used. All recordings had multiple z-slices within, above, and below the cell body permitting effective quantification of recordings with slight z-drift over hours of recording. For example, in Figure 1g, 14 z-slices were taken at 3 µm steps and only ∼5-6 of the z-slices included fluorescence from the oviDNb cell body. The length of each recording (mean of 75 min) varied depending on (1) the perceived health of the fly; (2) the chance of future egg-laying events (which were higher if the fly had already laid an egg); (3) the amount of z-drift; and (4) the quality of the agarose wheel, which sometimes visibly dried over a period of hours. The experimenter was blind to correlations between the neural signal and behavior during the vast majority (∼95%) of recordings. Flies were only excluded if a technical issue arose (e.g., errors synchronizing behavior with two-photon imaging or saline leaking from the holder). Only eggs with continuous two-photon imaging from 240 s before to 30 s after egg-deposition were analyzed.

For CsChrimson optogenetics experiments supporting a rise-to-threshold mechanism (Fig. 2f-h), two-photon imaging parameters were held relatively constant (mean framerate of 1.81 Hz, and two-photon laser power ∼10.5 mW). CsChrimson stimulation intensities were determined in pilot experiments. Periodic stimulation cycling 4 intensities was applied for 5 s every 2 min. The experimenter was blind to correlations between the neural signal and behavior during all these recordings.

For CsChrimson optogenetics experiments in Figure 2c, two-photon imaging data are only shown for 5 of the 9 flies, whereas the behavioral data are shown for all 9 flies. The 4 flies for whom we did not show imaging data had bleed-through artifacts in the GCaMP signal from the CsChrimson illumination LED because these data were collected prior to us optimizing the detection path for minimizing this artifact.

Two-photon imaging frames were motion corrected using custom scripts from a previous study^81^ or using CaImAn^84^. The ROIs for a cell body were drawn manually for each z-plane using the time-average of each z-plane. ROIs were drawn around the outer boundary of the cell body. The brighter of the two cell bodies in oviDN-SS1 was assigned to be oviDNb (Extended Data Fig. 6a). In a few cases, where the brighter cell was not obvious, ROIs encompassing both cell bodies were drawn and assigned to be oviDNb. For a given imaging volume timepoint, the individual pixel intensities in all the individual z-plane ROIs for a given cell were pooled and averaged, F_cell_(t). An identical average was done for a volume that appeared to be background, F_background_(t). Prior to calculating the ΔF/F, we subtracted the background from the cell, F_cell_actual_(t) = F_cell_(t) - F_background_(t). This eliminated non-cell-specific signal such as autofluorescence and constant detector background. This subtraction also made the ΔF/F robust to variations in the number of background pixels included in the ROIs drawn around the outside of a cell. ΔF/F was calculated using the formula, (F_cell_actual_(t) – F_0_(t)) / F_0_(t), where F_0_(t) was the running mean of F_cell_actual_(t) over a 20 min. window. The mean over a long timeframe was used to estimate a ‘baseline’, systematically, for the continuously fluctuating oviDN signal. A similar running mean ‘baseline’ estimate (albeit with a much shorter window) was used to quantify continuously fluctuating dopaminergic signals in mammals^85^. A ΔF/F of 0.35, for example, indicated that the fluorescence signal in the cell was 35% greater than the 20-min-mean signal in the cell. If the GCaMP7f fluorescence signal is linear with [Ca^2+^] in this range it would indicate that the [Ca^2+^] in the cell increased by 35% over the 20-min-mean [Ca^2+^] in the cell. All stated conclusions were robust to three different methodologies for calculating a ΔF/F, including methods where F_0_ remained constant. For CsChrimson experiments, F_0_(t) was the running mean of F_cell_actual_(t) over a 20 min. window after the 105 s post triggering CsChrimson stimulation was set to NaN. This very conservatively prevented any CsChrimson stimulations, or lingering effects, from artificially influencing F_0_(t).

Two-photon imaging-frame pulses, behavioral-camera frame triggers, and optogenetic LED triggers were all digitized at 10 kHz on a Digidata 1440A (Molecular Devices) and saved to a computer (Axoscope, Molecular Devices). To assign a timestamp to a volume scan, we identified the moment that the two-photon volume scan was half complete. To assign a timestamp to a behavioral camera frame, we used the beginning of the 16 ms camera exposure period. Calcium imaging were interpolated and behavioral data were down-sampled to a common 10 Hz array for all population analyses. Each 100 ms timepoint was assigned the calcium imaging and behavior data value from the closest previous respective timestamp (i.e., previous neighbor interpolation). A relatively large 100 ms time-base was chosen because faster sampling was unnecessary for the current analyses and would be computationally time-consuming given the 200+ hours of two-photon scanning collected. In the case of triggered averages, the zero point was the timestamp for the behavior camera frame with the behavior of interest or the frame with the onset of optogenetic stimulation. In the case of cross-correlations, the zero point was the timestamp of the first acquired two-photon volume.

### Electrophysiology

We used the same two-photon microscope for calcium imaging and for patch-clamp electrophysiology. The microscope was controlled by either Prairie View (Bruker) or µManager^86^ software. A 470 nm LED (pE-100, CoolLED) provided excitation through the objective to identify 2xEGFP or GCaMP7f positive neurons. An 850 nm LED coupled to a 400 µm wide lightguide (M850F2 and M28L01, Thorlabs) was focused on the fly’s head to illuminate cells for patch-clamping. Both LEDs were turned off when recording electrophysiology data. A 40x/0.80 NA objective (LUMPLFLN 40XW, Olympus) and CoolSnapEZ CCD camera (Photometrics) were used for patch-clamping.

Cell bodies were exposed by breaching the neural lamella and perineural sheath using gentle application of 0.5% collagenase IV (Worthington) in extracellular to a small 30 µm x 30 µm area containing the cell bodies of interest^82^. Collagenase was applied with a 4 to 6 µm tip micropipette with 8 to 80 mm Hg positive pressure at ∼30-32°C for ∼3 min. Once the cell bodies were exposed, the bath was returned to ∼19-21°C and flushed of collagenase.

Borosilicate glass (OD 1.5 mm, ID 0.86 mm, with filament) was pulled to create a 7 to 15 MΩ electrode with a 1 to 1.5 µm tip using a Model P-1000 Micropipette puller (Sutter Instruments). Intracellular saline^83^ was composed of 140 mM potassium-aspartate, 1 mM KCl, 10 mM HEPES, 1 mM EGTA, 0.5 mM Na_3_GTP, 4 mM MgATP, 13 mM biocytin hydrazide, and 20 µM Alexa-568–hydrazide-Na (ThermoFisher Scientific). The pH was adjusted to ∼7.3 with KOH and osmolarity was adjusted to ∼265 mOsm with water.

Electrophysiological signals were acquired using a MultiClamp 700B amplifier (Molecular Devices). The electrophysiological signals and behavioral camera triggers were digitized at 10 kHz via a Digidata 1440A and saved to a computer (Clampex, Molecular Devices). The oviDN or oviDN-like subtype (Extended Data Fig. 6a) that was recorded from was not distinguished. Recordings were made without current injection (except for current step protocols) and the reported membrane voltage (*Vm*) was corrected for a 13 mV junction potential^82^. Spikes were identified by highpass filtering the *Vm* and finding peaks above a threshold that were separated in time by > 1 ms. Parameters for peak detection were varied from recording to recording based on visual inspection of the data, in which the action potentials were clear. We calculated the spike rate by counting the number of spikes in every 5 s interval (at 0.1 ms steps), dividing by 5, and assigning that value to the middle of the 5 s interval. Spike rate and *Vm* were thus both measured at 0.1 ms intervals. Data were aligned and analyzed identically to calcium imaging. The resting *Vm* was considered the first stable *Vm* after breaking into the whole-cell configuration (Fig. 5d, Extended Data Fig. 12a). We calculated a *Vm* with spikes removed by discarding (converting to NaNs) 150 ms of data centered on the peak of each spike (Extended Data Fig. 12c).

Electrophysiological recordings for Extended Data Figure 12 were analyzed only if (1) the cell was stably recorded for more than 3 minutes; (2) the *Vm* was below −43 mV at rest with no large drift or rapid fluctuations in the V*m* that were non-physiological; (3) the fly walked for at least one wheel rotation; and (4) the cell spiked at least once. A total of 5 cells were rejected for not passing criteria 2, 3 and 4. 3 of these 5 were rejected for not passing criteria 2. A single cell was rejected for not passing criteria 3, indicating that flies were healthy in this preparation. A single cell passed the first 3 criteria but was rejected for not spiking (shown in Extended Data Fig. 12a). Cells that passed all 4 criteria were analyzed from the time the recording first stabilized to when the recording degraded or was terminated (mean = 41 min.).

Kir2.1* and Kir2.1*Mut flies were pre-treated as described for free behavior experiments, rather than as described for tethered experiments, so the transgene would be expressed as it was in free behavior. All recordings were done in vivo on the wheel. Recordings were analyzed if the *Vm* was below −43 mV at rest. Current step protocols were conducted with 5 pA increments with 1 s of current injection (Extended Data Fig. 15a).

### Texas Red Fill

100 mg/mL Texas Red (Dextran, Texas Red, 3000 MW, Lysine Fixable) (ThermoFisher Scientific) in patch-clamp intracellular saline (see above) lacking ATP, GTP, biocytin, or Alexa-568– hydrazide-Na was backfilled into a patch pipette. The pipette was positioned near the cell body (without any collagenase application) and 2 to 5 pulses of 10 V (2 ms duration) were applied using an SD9 stimulator (Grass Instruments). All fills and anatomy were done with flies on the wheel under the two-photon microscope (as in Calcium imaging, except using a 40x/0.80 NA objective (LUMPLFLN 40XW, Olympus) and a 590 to 650 nm bandpass filter (Chroma) to filter emitted light prior to entering a 2^nd^ GaAsP detector (Hamamatsu)).

### Substrate transition triggered averages during calcium imaging or electrophysiology

Substrate transitions were identified using the fly’s position on the wheel as described in ‘Design of egg-laying wheel…’. For these analyses, substrate transition *i* was eliminated if substrate transitions *i-1* and *i+1* occurred within 4 s of each other. This empirically prevented events where the fly rocked on the substrate boundary from counting as multiple transitions. Note that for all transition-triggered averages, if the fly were to have transitioned back to the original substrate, say, 20 s after the first transition, the data from 20 s onwards did not contribute to the post-transition average.

### Measurement of light power

All light power levels reported in this paper were measured with a PM100D Compact Power and Energy Console (Thorlabs) at the expected peak intensity of the light source. Lighting with area smaller than the sensor was divided by the estimated illuminated area, rather than the area of the sensor.

### Statistics

We used the two-sided Wilcoxon rank sum test to calculate all p-values. Error bars are standard error of the mean unless otherwise described.

For egg-laying choice fractions (like Fig. 3c) grey bars show the fraction of eggs laid on the lower sucrose option after all eggs from all flies are pooled. Error bars indicate the 95% confidence interval of this fraction calculated using the Clopper-Pearson method (‘exact’ binomial confidence interval). Individual dots indicate individual flies.

P-values in the main text compare the number of trials with (or without) events in two separate groups. For a single group, trials with an event are treated as 1 and trials without an event are treated as 0. Then the two groups (each a set of 0 and 1) are compared using the two-sided Wilcoxon rank sum test (p-values calculated using two-sided Fisher’s exact test are similar and similarly significant).

For all experiments, no data were excluded unless explicitly stated, and no statistical method was used to choose sample size.

### Data Analysis Software

All data analyses and instrument control were done using MATLAB (MathWorks) unless otherwise specified.

## Acknowledgements

We thank the Rockefeller University Precision Instrumentation Technologies facility for access to fabrication equipment; the Janelia FlyLight team for generating images of oviDN-GAL4 (in Fig. 1f and Extended Data Fig. 5b) and oviDN-SS1 MCFO (Fig. 1e); the Bloomington Drosophila Stock Center (NIH P400D018537) for various fly stocks; Andrew Siliciano and Vanessa Ruta for GtACR1 effector flies; Mariana Wolfer for flies expressing GCaMP in eggs; Massimo Scanziani for pCAG-Kir2.1Mut-T2A-tdTomato and pCAG-Kir2.1-T2A-tdTomato plasmids (Addgene plasmid # 60644 and 60598); Gerald Rubin for pJFRC81-10XUAS-IVS-Syn21-GFP-p10 plasmid (Addgene plasmid # 36432); Meishel DeSouto for template fly drawing that was modified and used throughout the manuscript; Zikun Wang for help developing the first iteration of the fly tracking setup; Kris Fonselius and Sam Cohen for involvement in developing tools for computer-assisted manual annotation of egg-deposition events in free behavior; Joseph Varikooty, Jonathan Hirokawa and Itzel Ishida for ideas and help developing the wheel; Jazz Weisman for sharing his design for delivery of optogenetic stimulation light; and Cheng Lyu and Jonathan Green for two-photon and electrophysiology discussions. Research reported in this publication was supported by a Brain Initiative grant from the National Institute of Neurological Disorders and Stroke (R01NS121904) to G.M. and Leon Levy Foundation fellowship and the Kavli Neural Systems Institute grant to V.V.. G.M. is a Howard Hughes Medical Institute Investigator.

## Author Contributions

V.V. and G.M. conceived the initial study and wrote the manuscript. V.V., with input from G.M., designed the experiments, performed the experiments, analyzed the data, interpreted the results, and decided on new experiments. F.W., K.W., and B.J.D. provided oviDN genetic driver lines and their anatomical characterization, shared preliminary data on the oviDNs, and provided helpful feedback on experiments and the manuscript. A.C. and A.A. created the Kir2.1* and Kir2.1*Mut flies. H.A. developed code for computer-assisted manual annotation of egg-deposition events in free behavior.

The authors declare no competing interests.

## Author Information

Correspondence and requests for materials should be addressed to

## Data and Resources Availability

Datasets, code and technical documents (e.g., CAD files and plasmid maps) used in this study are available from the corresponding authors on request and will be made publicly available before or at time of publication.

## References

1. Masset, P. & Kepecs, A. Categorical Decisions. in Encyclopedia of Computational Neuroscience (eds. Jaeger, D. & Jung, R.) 1–5 (Springer, 2013). doi:10.1007/978-1-4614-7320-6_310-1.

2. Gold, J. I. & Shadlen, M. N. The Neural Basis of Decision Making. Annu. Rev. Neurosci. 30, 535–574 (2007).

3. Hanks, T. D. & Summerfield, C. Perceptual Decision Making in Rodents, Monkeys, and Humans. Neuron 93, 15–31 (2017).

4. Shadlen, M. N. & Newsome, W. T. Motion perception: seeing and deciding. PNAS 93, 628–633 (1996).

5. Daw, N. D., O’Doherty, J. P., Dayan, P., Seymour, B. & Dolan, R. J. Cortical substrates for exploratory decisions in humans. Nature 441, 876–879 (2006).

6. Hayden, B. Y., Pearson, J. M. & Platt, M. L. Neuronal basis of sequential foraging decisions in a patchy environment. Nat Neurosci 14, 933–939 (2011).

7. Kolling, N., Behrens, T. E. J., Mars, R. B. & Rushworth, M. F. S. Neural Mechanisms of Foraging. Science 336, 95–98 (2012).

8. Hayden, B. Y. Economic choice: the foraging perspective. Current Opinion in Behavioral Sciences 24, 1–6 (2018).

9. Yang, C., Belawat, P., Hafen, E., Jan, L. Y. & Jan, Y.-N. Drosophila Egg-Laying Site Selection as a System to Study Simple Decision-Making Processes. Science 319, 1679–1683 (2008).

10. Yang, C.-H., He, R. & Stern, U. Behavioral and Circuit Basis of Sucrose Rejection by Drosophila Females in a Simple Decision-Making Task. J. Neurosci. 35, 1396–1410 (2015).

11. Wang, F. et al. Neural circuitry linking mating and egg laying in Drosophila females. Nature 579, 101–105 (2020).

12. Cury, K. M., Prud’homme, B. & Gompel, N. A short guide to insect oviposition: when, where and how to lay an egg. Journal of Neurogenetics 33, 75–89 (2019).

13. Karageorgi, M. et al. Evolution of Multiple Sensory Systems Drives Novel Egg-Laying Behavior in the Fruit Pest Drosophila suzukii. Current Biology 27, 847–853 (2017).

14. Zhu, E. Y., Guntur, A. R., He, R., Stern, U. & Yang, C.-H. Egg-laying demand induces aversion of UV light in Drosophila females. Curr Biol 24, 2797–2804 (2014).

15. Stensmyr, M. C. et al. A conserved dedicated olfactory circuit for detecting harmful microbes in Drosophila. Cell 151, 1345–1357 (2012).

16. Liu, W. et al. Enterococci Mediate the Oviposition Preference of Drosophila melanogaster through Sucrose Catabolism. Scientific Reports 7, 13420 (2017).

17. Azanchi, R., Kaun, K. R. & Heberlein, U. Competing dopamine neurons drive oviposition choice for ethanol in Drosophila. Proc Natl Acad Sci U S A 110, 21153–21158 (2013).

18. Kacsoh, B. Z., Lynch, Z. R., Mortimer, N. T. & Schlenke, T. A. Fruit Flies Medicate Offspring After Seeing Parasites. Science 339, 947–950 (2013).

19. Joseph, R. M., Devineni, A. V., King, I. F. G. & Heberlein, U. Oviposition preference for and positional avoidance of acetic acid provide a model for competing behavioral drives in Drosophila. PNAS 106, 11352–11357 (2009).

20. Hussain, A. et al. Ionotropic Chemosensory Receptors Mediate the Taste and Smell of Polyamines. PLOS Biology 14, e1002454 (2016).

21. Dweck, H. K. M. et al. Olfactory Preference for Egg Laying on Citrus Substrates in Drosophila. Current Biology 23, 2472–2480 (2013).

22. Joseph, R. M. & Heberlein, U. Tissue-Specific Activation of a Single Gustatory Receptor Produces Opposing Behavioral Responses in Drosophila. Genetics 192, 521–532 (2012).

23. Bräcker, L. B. et al. Quantitative and Discrete Evolutionary Changes in the Egg-Laying Behavior of Single Drosophila Females. Front. Behav. Neurosci. 13, (2019).

24. Cury, K. M. & Axel, R. Decisions in an Innate Behavioral Sequence. bioRxiv 2021.04.03.438315 (2021) doi:10.1101/2021.04.03.438315.

25. Oliveira-Ferreira, C., Gaspar, M. & Vasconcelos, M. L. Neuronal control of suppression, initiation and completion of egg deposition in Drosophila melanogaster. 2021.08.23.457359 https://www.biorxiv.org/content/10.1101/2021.08.23.457359v1 (2021) doi:10.1101/2021.08.23.457359.

26. Heifetz, Y., Yu, J. & Wolfner, M. F. Ovulation Triggers Activation of Drosophila Oocytes. Developmental Biology 234, 416–424 (2001).

27. Tian, L. et al. Imaging neural activity in worms, flies and mice with improved GCaMP calcium indicators. Nat Methods 6, 875–881 (2009).

28. Kaneuchi, T. et al. Calcium waves occur as Drosophila oocytes activate. PNAS 112, 791–796 (2015).

29. Scheffer, L. K. et al. A connectome and analysis of the adult Drosophila central brain. eLife 9, e57443 (2020).

30. Dana, H. et al. High-performance calcium sensors for imaging activity in neuronal populations and microcompartments. Nature Methods 16, 649–657 (2019).

31. Berridge, M. J., Bootman, M. D. & Lipp, P. Calcium - a life and death signal. Nature 395, 645–648 (1998).

32. Klapoetke, N. C. et al. Independent optical excitation of distinct neural populations. Nature Methods 11, 338–346 (2014).

33. Yang, H. H. et al. Subcellular Imaging of Voltage and Calcium Signals Reveals Neural Processing In Vivo. Cell 166, 245–257 (2016).

34. Baines, R. A., Uhler, J. P., Thompson, A., Sweeney, S. T. & Bate, M. Altered Electrical Properties in DrosophilaNeurons Developing without Synaptic Transmission. J. Neurosci. 21, 1523–1531 (2001).

35. Mohammad, F. et al. Optogenetic inhibition of behavior with anion channelrhodopsins. Nature Methods 14, 271–274 (2017).

36. Xue, M., Atallah, B. V. & Scanziani, M. Equalizing excitation–inhibition ratios across visual cortical neurons. Nature 511, 596–600 (2014).

37. Harrell, E. R., Pimentel, D. & Miesenböck, G. Changes in Presynaptic Gene Expression during Homeostatic Compensation at a Central Synapse. J. Neurosci. 41, 3054–3067 (2021).

38. Flavell, S. W. & Greenberg, M. E. Signaling mechanisms linking neuronal activity to gene expression and plasticity of the nervous system. Annu Rev Neurosci 31, 563–590 (2008).

39. Ali, F. & Kwan, A. C. Interpreting in vivo calcium signals from neuronal cell bodies, axons, and dendrites: a review. Neurophotonics 7, 011402 (2020).

40. Dodge, F. A. & Rahamimoff, R. Co-operative action a calcium ions in transmitter release at the neuromuscular junction. J Physiol 193, 419–432 (1967).

41. Rg, O., Pm, D. & Sp, K. A supramodal accumulation-to-bound signal that determines perceptual decisions in humans. Nat Neurosci 15, 1729–1735 (2012).

42. Kelly, S. P. & O’Connell, R. G. Internal and External Influences on the Rate of Sensory Evidence Accumulation in the Human Brain. J. Neurosci. 33, 19434–19441 (2013).

43. Hanes, D. P. & Schall, J. D. Neural Control of Voluntary Movement Initiation. Science 274, 427–430 (1996).

44. Roitman, J. D. & Shadlen, M. N. Response of neurons in the lateral intraparietal area during a combined visual discrimination reaction time task. J Neurosci 22, 9475–9489 (2002).

45. Maimon, G. & Assad, J. A. A cognitive signal for the proactive timing of action in macaque LIP. Nature Neuroscience 9, 948–955 (2006).

46. Quintana, J. & Fuster, J. M. From perception to action: temporal integrative functions of prefrontal and parietal neurons. Cereb Cortex 9, 213–221 (1999).

47. Guo, Z. V. et al. Flow of Cortical Activity Underlying a Tactile Decision in Mice. Neuron 81, 179–194 (2014).

48. Hanks, T. D. et al. Distinct relationships of parietal and prefrontal cortices to evidence accumulation. Nature 520, 220–223 (2015).

49. Chen, T.-W., Li, N., Daie, K. & Svoboda, K. A Map of Anticipatory Activity in Mouse Motor Cortex. Neuron 94, 866–879.e4 (2017).

50. Evans, D. A. et al. A synaptic threshold mechanism for computing escape decisions. Nature 558, 590–594 (2018).

51. Steinmetz, N. A., Zatka-Haas, P., Carandini, M. & Harris, K. D. Distributed coding of choice, action and engagement across the mouse brain. Nature 576, 266–273 (2019).

52. Chabrol, F. P., Blot, A. & Mrsic-Flogel, T. D. Cerebellar Contribution to Preparatory Activity in Motor Neocortex. Neuron 103, 506–519.e4 (2019).

53. Mu, Y. et al. Glia Accumulate Evidence that Actions Are Futile and Suppress Unsuccessful Behavior. Cell 178, 27–43.e19 (2019).

54. Bahl, A. & Engert, F. Neural circuits for evidence accumulation and decision making in larval zebrafish. Nat Neurosci 23, 94–102 (2020).

55. Dragomir, E. I., Štih, V. & Portugues, R. Evidence accumulation during a sensorimotor decision task revealed by whole-brain imaging. Nature Neuroscience 23, 85–93 (2020).

56. von Reyn, C. R. et al. A spike-timing mechanism for action selection. Nature Neuroscience 17, 962–970 (2014).

57. DasGupta, S., Ferreira, C. H. & Miesenböck, G. FoxP influences the speed and accuracy of a perceptual decision in Drosophila. Science 344, 901–904 (2014).

58. Groschner, L. N., Chan Wah Hak, L., Bogacz, R., DasGupta, S. & Miesenböck, G. Dendritic Integration of Sensory Evidence in Perceptual Decision-Making. Cell 173, 894–905.e13 (2018).

59. McKellar, C. E. et al. Threshold-Based Ordering of Sequential Actions during Drosophila Courtship. Current Biology 29, 426–434.e6 (2019).

60. Fotowat, H., Harrison, R. R. & Gabbiani, F. Multiplexing of Motor Information in the Discharge of a Collision Detecting Neuron during Escape Behaviors. Neuron 69, 147–158 (2011).

61. Mann, K., Gordon, M. D. & Scott, K. A Pair of Interneurons Influences the Choice between Feeding and Locomotion in Drosophila. Neuron 79, 754–765 (2013).

62. Kim, I. S. & Dickinson, M. H. Idiothetic Path Integration in the Fruit Fly Drosophila melanogaster. Current Biology 27, 2227–2238.e3 (2017).

63. Corfas, R. A., Sharma, T. & Dickinson, M. H. Diverse Food-Sensing Neurons Trigger Idiothetic Local Search in Drosophila. Curr Biol 29, 1660–1668.e4 (2019).

64. Ofstad, T. A., Zuker, C. S. & Reiser, M. B. Visual place learning in Drosophila melanogaster. Nature 474, 204–207 (2011).

65. Simon, J. C. & Dickinson, M. H. A New Chamber for Studying the Behavior of Drosophila. PLOS ONE 5, e8793 (2010).

66. Robertson, F. W., Sang, J. H. & Hogben, L. T. The ecological determinants of population growth in a Drosophila culture. I. Fecundity of adult flies. Proceedings of the Royal Society of London. Series B - Biological Sciences 132, 258–277 (1944).

67. Mathis, A. et al. DeepLabCut: markerless pose estimation of user-defined body parts with deep learning. Nature Neuroscience 21, 1281–1289 (2018).

68. Gou, B., Zhu, E., He, R., Stern, U. & Yang, C.-H. High Throughput Assay to Examine Egg-Laying Preferences of Individual Drosophila melanogaster. J Vis Exp (2016) doi:10.3791/53716.

69. Feiler, R., Harris, W. A., Kirschfeld, K., Wehrhahn, C. & Zuker, C. S. Targeted misexpression of a Drosophila opsin gene leads to altered visual function. Nature 333, 737–741 (1988).

70. Yamaguchi, S., Desplan, C. & Heisenberg, M. Contribution of photoreceptor subtypes to spectral wavelength preference in Drosophila. Proc Natl Acad Sci U S A 107, 5634–5639 (2010).

71. Sharkey, C. R., Blanco, J., Leibowitz, M. M., Pinto-Benito, D. & Wardill, T. J. The spectral sensitivity of Drosophila photoreceptors. Scientific Reports 10, 18242 (2020).

72. Branson, K., Robie, A., Bender, J., Perona, P. & Dickinson, M. High-throughput Ethomics in Large Groups of Drosophila. Nat Methods 6, 451–457 (2009).

73. McGuire, S. E., Le, P. T., Osborn, A. J., Matsumoto, K. & Davis, R. L. Spatiotemporal Rescue of Memory Dysfunction in Drosophila. Science 302, 1765–1768 (2003).

74. Mauss, A. S., Busch, C. & Borst, A. Optogenetic Neuronal Silencing in Drosophila during Visual Processing. Scientific Reports 7, 13823 (2017).

75. Johns, D. C., Marx, R., Mains, R. E., O’Rourke, B. & Marbán, E. Inducible Genetic Suppression of Neuronal Excitability. J. Neurosci. 19, 1691–1697 (1999).

76. Hardie, R. C. et al. Calcium Influx via TRP Channels Is Required to Maintain PIP2 Levels in Drosophila Photoreceptors. Neuron 30, 149–159 (2001).

77. Pfeiffer, B. D., Truman, J. W. & Rubin, G. M. Using translational enhancers to increase transgene expression in Drosophila. Proc Natl Acad Sci U S A 109, 6626–6631 (2012).

78. Kim, A. J., Fenk, L. M., Lyu, C. & Maimon, G. Quantitative Predictions Orchestrate Visual Signaling in Drosophila. Cell 168, 280–294.e12 (2017).

79. Okubo, T. S., Patella, P., D’Alessandro, I. & Wilson, R. I. A Neural Network for Wind-Guided Compass Navigation. Neuron 107, 924–940.e18 (2020).

80. Seelig, J. D. et al. Two-photon calcium imaging from head-fixed Drosophila during optomotor walking behavior. Nature Methods 7, 535–540 (2010).

81. Green, J. et al. A neural circuit architecture for angular integration in Drosophila. Nature 546, 101–106 (2017).

82. Maimon, G., Straw, A. D. & Dickinson, M. H. Active flight increases the gain of visual motion processing in Drosophila. Nature Neuroscience 13, 393–399 (2010).

83. Wilson, R. I. & Laurent, G. Role of GABAergic Inhibition in Shaping Odor-Evoked Spatiotemporal Patterns in the Drosophila Antennal Lobe. J. Neurosci. 25, 9069–9079 (2005).

84. Giovannucci, A. et al. CaImAn an open source tool for scalable calcium imaging data analysis. eLife 8, e38173 (2019).

85. Hamilos, A. E. et al. Dynamic dopaminergic activity controls the timing of self-timed movement. bioRxiv 2020.05.13.094904 (2020) doi:10.1101/2020.05.13.094904.

86. Edelstein, A., Amodaj, N., Hoover, K., Vale, R. & Stuurman, N. Computer Control of Microscopes Using µManager. Current Protocols in Molecular Biology 92, 14.20.1–14.20.17 (2010).

